# Widespread evidence for plasticity and recent evolution of plasticity in the breeding phenology of Finnish birds

**DOI:** 10.64898/2026.06.05.730291

**Authors:** M.H Hällfors, A Lehikoinen, A.B Phillimore

## Abstract

Phenological shifts under climate change often arise through phenotypic plasticity and, where this is insufficient to track shifts in optimum timing, genetic adaptation may also play a role. Understanding the contributions of these two processes is critical for predicting species persistence in a changing climate. While many species show phenological plasticity, we know surprisingly little about the contributions that genetic adaptation of the plasticity reaction norm elevation (timing in the mean environment) and slope (shift in timing as a response to temperature) make to phenological shifts. With the aim of disentangling plasticity from adaptation in temperature–phenology reaction norms, we applied a statistical approach to long-term first egg-laying data from 44 Finnish bird species represented by 69 populations spanning six decades. Applying phylogenetic meta-analysis to parameter estimates obtained from the individual time series, we estimated average plasticity and adaptation effect sizes and tested whether migratory strategy, generation length, and mean laying-date explained among-species variation. Egg-laying phenology was strongly plastic, advancing by 2.5 days °C□¹. We found no evidence for a steeper reaction norm between 5-year periods versus within them, consistent with no adaptation of the reaction norm elevation. However, we detected a significant steepening of slopes over time (−0.04 days °C□¹ year□¹), consistent with plasticity across the whole study area increasing from -2.5 to -5.1 days °C□¹ and in the northernmost area (−0.07 days °C□¹ year□¹) from -2 to -6.5 days °C□¹ over the 64-year study period. Trait analyses revealed no significant effect of migratory strategy, generation length, nor mean phenology on adaptation. We show that plasticity enables substantial short-term tracking of warming accompanied by noteworthy evidence consistent with widespread evolution of. Our approach demonstrates how observational data can help reveal evolutionary signals, offering a tool for improved understanding of the processes that underpin phenological responses.

## 1. Introduction

In response to climate change, species must either adjust to changing conditions or shift their geographical distributions toward emerging favorable areas (Thurman et al., 2020). Two mechanisms that allow species to adjust to new conditions where they currently occur are (i) adaptive phenotypic plasticity, whereby a genotype is able to produce different traits in different environments that tracks (or partially tracks) a shift in the optimum (Ghalambor et al., 2007) and (ii) evolutionary adaptation involving natural selection of heritable traits that enhance fitness under changing conditions (Holt, 1990). Where plasticity is insufficient for the trait value to track the shift in the optimum (Chevin et al. 2010) or reaches its limit (Gerlich et al., 2025; Iler et al., 2013; Schmidt et al., 2023), populations may lag behind the optimum (Radchuk et al., 2019) and experience directional selection (Alberto et al., 2013; Chevin et al., 2010; Duputié et al., 2015; Gienapp et al., 2008; Van Buskirk et al., 2012), potentially leading to adaptation of the reaction norm elevation (i.e., phenological timing in the average environment) or slope (i.e., sensitivity of the phenotype to the environment).

We have substantial evidence that phenotypic plasticity makes an important contribution to phenotypic change in response to climate change, owing to its rapid expression and relative tractability in both field and laboratory studies (Merilä, 2012; Merilä & Hendry, 2014). In comparison, genetic adaptation builds up over multiple generations, and there are a scarcity of empirical studies quantifying the contribution that adaptation to climate change makes to phenotypic evolution over time (Hansen et al., 2012; Merilä, 2012; Merilä & Hendry, 2014; Ramakers et al., 2019). Importantly, where the combination of plasticity and adaptation (of reaction norm elevation or slope) is insufficient to track the optimum phenotype (Chevin et al., 2010; Gao et al., 2018; Radchuk et al., 2019, 2026), this may result in population decline and eventually extinction, even leading to reduced stability of the ecosystem (Rodrigues et al., 2025; Sgrò et al., 2011). Relying solely on plasticity may not be enough for populations to cope under continued rapid climate change (Duputié et al., 2015), and understanding the extent to which populations can evolve under current change could substantially improve our ability to predict population fates. Thus, discerning the relative roles that plasticity and adaptive evolution have played to date provides a guide both to how populations are responding currently and how they may continue to respond in the future. However, the paucity of empirical studies quantifying genetic adaptation limits our ability to draw general conclusions on the relative roles of plasticity and adaptation (including the evolution of plasticity), and how and why their contributions vary among species.

Demonstrating adaptive evolution in response to anthropogenic change is challenging (Hansen et al., 2012). Powerful methods that can be applied to study genetic change and adaptation of traits over time include (i) resurrection studies, which typically involves the comparison of fitness between contemporary and resurrected individuals under current conditions (Franks et al. 2014, 2018b, Merilä and Hendry 2014; Thomann et al. 2015), (ii) common garden experiments replicated in time, wherein the mean phenotypes of populations sampled at different points in time can be compared whilst controlling for environmental effects (Bradshaw & Holzapfel, 2001; Helm et al., 2019; van Asch et al., 2013), and (iii) quantitative genetic studies using multi-generation pedigrees, whereby a change in trait breeding value over time or in relation to a driver can be compared with the change expected under genetic drift (Bonnet et al., 2019). However, only a handful of species are amenable to such experiments or have been subject to the intensive long-term study required to build a pedigree (Charmantier & Gienapp, 2014; Ramakers et al., 2019; van Asch et al., 2013). Against this backdrop, approaches that enable estimation of genetic adaptation of traits from the wealth of long-term observational phenotypic data from various monitoring programmes would greatly extend the breadth of taxa and regions for which we could gain insights into genetic adaptation.

Here, we present an approach that uses long-term monitoring data on population phenotypes coupled with temperature data and seeks to decompose the response into (i) an estimate of plasticity and (ii) estimates of the contributions of adaptation. An important starting point for our approach is to focus on a trait that likely is under selection and for which the optimum is hypothesized to be temperature sensitive (Merilä & Hendry, 2014). Reproductive phenology in high latitude environments satisfies both criteria. The mean timing and plasticity of phenology exhibited by a population has evolved as a response to the seasonal conditions experienced, with fitness consequences of misaligned breeding timing potentially being substantial (Radchuk et al., 2019, 2026). To test for a signature of genetic adaptation in phenology, we modify a correlation-based approach that has been used to estimate *local adaptation of a trait to temperature over space* (LAS; Phillimore et al., 2010, 2012), to function in a purely temporal setting and estimate *adaptation of a trait to temperature over time* (AT). The LAS approach is readily applicable to observational data for any species for which sufficient replication of population-level observations of phenotypic traits exists, such as for citizen science phenological data, without requiring direct information on individual fitness (Phillimore et al., 2012). The premise of the LAS approach is that, assuming that plasticity is a constant and the thermal cue has been identified, the population reaction norm (for a trait regressed on the temperature cue) estimated over short-term periods will be mainly attributable to plasticity, whereas the population reaction norm estimated over space (between populations) may capture a contribution of (local) adaptation in addition to plasticity (Fig. 1a and b; Phillimore et al. 2010). In transposing the LAS approach to a purely temporal axis (AT), we retain the assumption that, over short-term periods of time, the population reaction norm provides an estimate of population plasticity (Fig. 1a and c), whereas across longer time periods (multi-year averages; Fig. 1c-g), the reaction norm may in addition to plasticity, also capture contributions of adaptation over time. Under the *plasticity only hypothesis* for AT, the short- and long-term phenological response is the same (Fig. 1c). Given a directional trend in temperature, where the environmental sensitivity of the optimum (*B*, sensu Chevin et al. (2010)) exceeds the plastic reaction norm (b) and therefore the population phenotypic response is lagging behind the optimum, adaptation may shift the phenotype toward the optimum. Under this *elevation adaptation hypothesis*, the phenotype-temperature slope across longer time periods is predicted to be steeper than that within short-term periods (Fig. 1d). Alternatively, or in addition, under the *slope adaptation hypothesis*, adaptation may lead to a steepening of the plastic reaction norm slope across the time-series (Fig. 1g).

**Figure 1.**
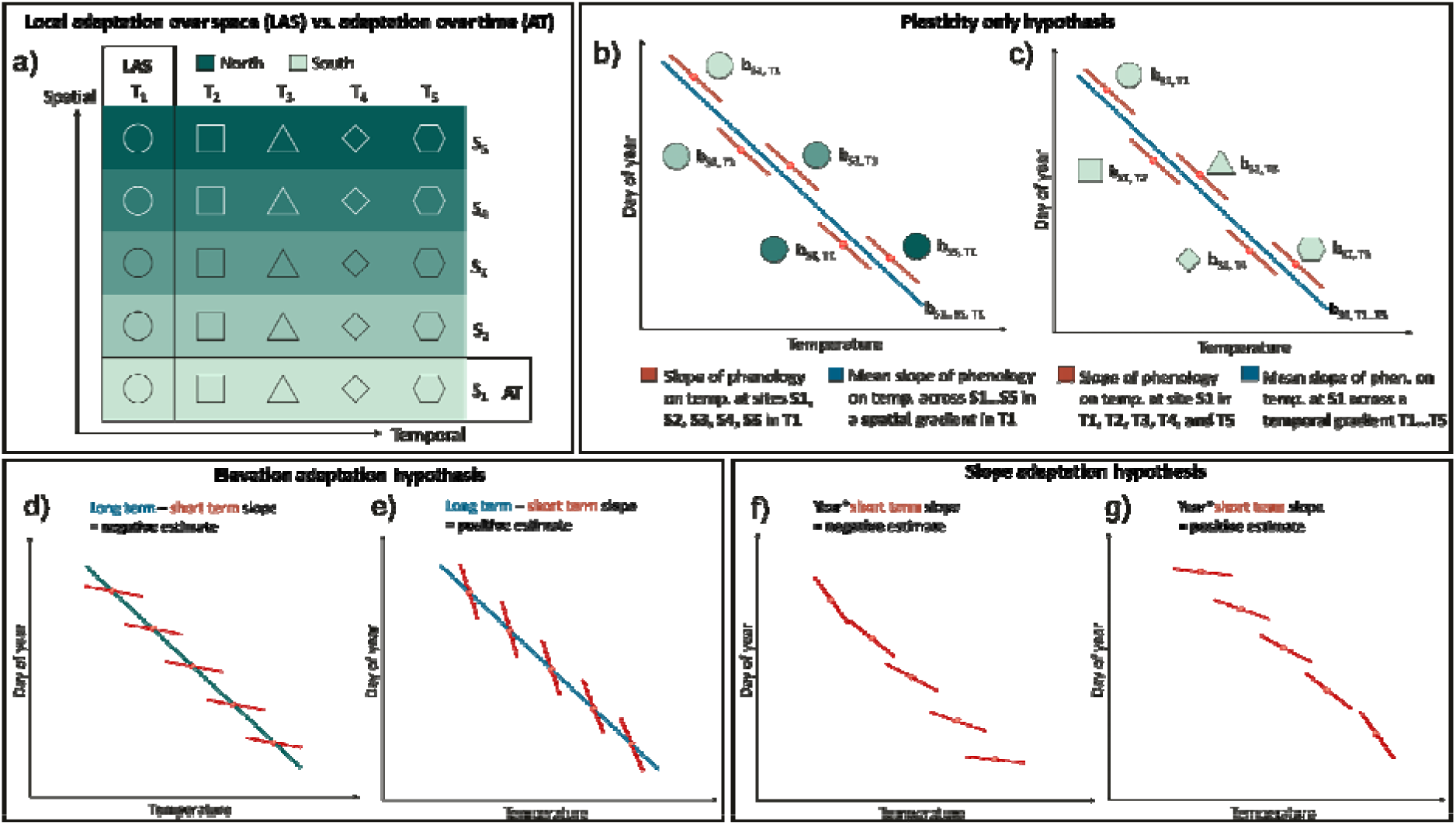
Schematic representation of our approach to disentangle adaptive evolution from plastic responses based on observational data and the regression of phenology on a temperature cue. Panel a-c shows the difference between the local adaptation over space approach (LAS; Phillimore et al., 2010, 2012) and our deployment of the same statistical approach to capture adaptation over time (AT). Panel b depicts the LAS approach (first column in panel a) and a case where there is no local adaptation, and the spatial slope and a temporal slope estimated across a short temporal gradient (T_1_). To transfer the LAS approach and function in a temporal setting (AT; bottom row in panel a), we define local populations as a set of individuals occurring in the same geographical region and group observations done during a short period of time (e.g., a five-year period). In AT, the subject is this regionally defined population under a defined temporal period (population S_1_ during each time period T_1_-T_5_). Where T corresponds to a multi-year period that is short enough to include only a small number of generations, the slope of phenology regressed on temperature is assumed to provide an estimate of mean population phenological plasticity without an extensive impact of evolution within T. Panel c depicts the AT approach under the *plasticity-only hypothesis*, where the estimate of short-term plasticity (estimated within each short-term period, T_1_-T_5_, and averaged across them) is equal to the long-term slope estimated across the mean temperatures for each time period (mean temperature in T_1_-T_5_). Panels d-g depict scenarios where the long-term and short-term slopes differ. Under the *elevation adaptation hypothesis*, the average short-term response across populations is consistently shallower (panel d) or steeper (panel e) than the long-term response and would be consistent with co-gradient and counter-gradient adaptation of the population reaction norm elevation, respectively. Under the *slope adaptation hypothesis,* the short-term plastic response varies among the time periods (panels f-g), caused by evolution of the reaction norm slope.

In this study, we focus on breeding phenology in birds, because of its strong relationship to fitness and its sensitivity to thermal conditions (Charmantier & Gienapp, 2014; McLean et al., 2022). Populations and species may vary both in their plasticity and in potential signals of evolution of the reaction norms, and some of this variation may be explained based on differences in ecology and life history. For instance, migratory strategy (residents, short-distance migrants, or long-distance migrants) has previously been identified as a predictor of bird species’ tendency to shift their phenology (Hällfors et al., 2020; Lehikoinen et al., 2019; Saino et al., 2011), with resident and short-distance migratory species tending to arrive earlier and thus breed earlier in the season than long-distance migrants (Kluen et al., 2017). This may also place long-distant migrants under stronger selection for earlier breeding, and such evolutionary advancements have been reported for pied flycatchers (Helm et al., 2019; Lamers et al., 2023). Similarly, the average phenology of a species (early breeder versus late breeder) may affect the selection pressure that the species is exposed to, as seasonal transitions in temperature are more pronounced in the early spring compared to later spring and early summer, often including occasional cold spells. Therefore, we hypothesise that breeding phenology may be under stronger selection for populations breeding early in the season. Generation time is connected to a species’ “pace of life” (Healy et al., 2019) and therefore more short-lived species may be expected to evolve more rapidly to a changing environment (Compagnoni et al., 2021; Franks et al., 2007; Urban et al., 2023).

To capture bird breeding phenology, we used bird first egg-laying date (FELD) collected through citizen science over 64 years in Finland for 44 species sampled across four latitudinal bioclimatic zones. We then estimate if and when each population is most sensitive to spring temperatures. Even with time-series of multiple decades, we expect the power to detect a population-specific signal of adaptation (Fig. 1) to be very low. Therefore, we adopt a phylogenetic meta-analytic approach with the aim of estimating average effect sizes across populations (cf. Gerlich et al., 2025). We have four main aims. The first three are focused on quantifying the mean effects size of population plasticity (aim 1), and adaptation of the population reaction norm elevation (aim 2, *elevation adaptation hypothesis*, Fig. 1d-e) and reaction norm slope (aim 3, *slope adaptation hypothesis*, Fig. 1f-g). Our fourth aim is to examine the ability of traits (migratory strategy, early versus late breeding strategy, and generation time) to explain among species variation in effect sizes.

## 2. Materials and methods

### 2.1. Bird nesting data and estimation of FELD

We used data from the Finnish nest-card scheme collected during 1961–2025 (described in Kluen et al. 2017). Observations of breeding phases in bird nests are done by trained volunteer ornithologists in a standardized manner. Nests are usually visited repeatedly during the breeding season and observations on the presence of eggs, clutch size, and the development stages of nestlings are recorded. The data has broad spatial, temporal, and taxonomic coverage (whole of Finland, starting in 1901, 236 species out of all 250 breeding species, 249 807 recorded nests; Fig. S1) but with variation in records across space, time, and taxa.

In most nest cards, the exact first egg-laying date (date as in day of year; hereafter FELD) is unknown, as the recorder rarely visits the nest on the day the first egg is laid. Thus, we estimated FELD using a set of 26 guidelines that are informed by biological breeding parameters (as described by Kluen et al., (2017)). By using these guidelines, we were also able to estimate the likely accuracy of the estimated FELD. In general, the accuracy is higher when nest cards include more information (i.e., more visits) and visits closer to the FELD. If different guideline gave rise to differing estimates with differing accuracy levels, the estimate associated with the highest accuracy was used. The accuracy levels were converted into measurement error variance around the estimated day of hatching (Text S1), which we used as weights in subsequent models.

To apply the adaptation over time -approach (AT), we distinguished between not only populations in space (to discern between separate regions across the large study area) but, importantly, also defined temporally distinct groups of observations within these populations across the study period (cf. Fig. 1a and c). This means that for a single population we compare the short-term reaction norm within distinct time periods (5-years, estimating plasticity) to the reaction norm estimated between these time periods. Our decision to consider 5-year periods was motivated by our aim of maximizing power to estimate both the within and between-period slopes.

We selected data from 1961 onwards, to coincide with the availability of daily weather data (described in section 2.2). After FELD estimation and removal of records prior to 1961, the complete data contained 231 596 nest records of 236 species visited during 1961–2025 (Fig. S1a-b). As these data are based on citizen science recording, we are unable to follow spatially high-resolution local populations across time. Instead, we aggregate bioclimatic zone-specific (Ahti et al. 1968) observations of individual species which, within each bioclimatic zone, had been recorded during at least 20 years with at least five or more records during each of those years, and with these years being distributed across at least five of the 13 short-term periods, and treat them as regional populations. This left us with 77 species and 141 populations: 30 species in the hemiboreal zone, 72 in the southern boreal zone, 30 in the middle boreal zone, and nine species in the northern boreal zone (Fig. S1 c-d; data available in Hällfors et al., 2026).

### 2.2. Temperature data

We obtained daily temperature data at a 10□×□10□km resolution across Finland (interpolation methods described in Aalto et al., 2016) and estimated the average daily values across bioclimatic zones, allowing us to focus solely on the temporal (rather than spatiotemporal) impact of temperature on the phenology of each species in each bioclimatic zone. We also used these data to estimate temperature change during cue search windows (day of the year 60-170; see Section 2.4) across the study period in all potential sites across Finland, in the observation sites of bird breeding, and within the identified population-specific cue windows.

### 2.3. Trait and phylogenetic data

We obtained trait data describing migratory strategy and generation length based on literature (Cramp et al., 1994; Lehikoinen & Virkkala, 2016; Solonen, 1985; Valkama et al., 2014) and estimated average day of year when the species breeds from the same data as used for the main models. To quantify pace of life we used generation time, measured as mean longevity in years.

We extracted 100 phylogenies of the analyzed species using Birdtree.org (Jetz et al., 2012, 2014) and generated a consensus tree using the *consensus* function in the R package *ape* (Paradis et al., 2004).

### 2.4. Sliding window approach for cue window identification and removing populations with uncertain cue windows

Before estimating temperature-phenology reaction norms, it is important to identify (i) the environmental cues that a population’s FELD is most responsive to and (ii) whether FELD is temperature sensitive. The approach is described in Text S2.

Even under a null scenario where a species is not responsive to a thermal cue during the spring, the sliding window approach will still find a window that yields the lowest AIC (Bailey & Pol, 2016). To minimise the potential for downstream analyses to include populations for which there is little evidence for a relationship between temperature and phenology, we adopted two approaches to identify such populations and exclude them (described in Text S3). Adding theses two filters resulted in retention of 69 populations represented by 44 species (49% of our original number of populations in the data set). For the analyses, we thus ended up with 13 species in the hemiboreal zone, 37 in the southern boreal zone, 14 in the middle boreal zone, and five in the northern boreal zone (Fig. 2; Fig. S1 e-f). As a robustness test, we also applied an alternative approach (ii), using a more conservative threshold of the second approach, which retained 53 populations of 35 species (37% of populations).

**Figure 2.**
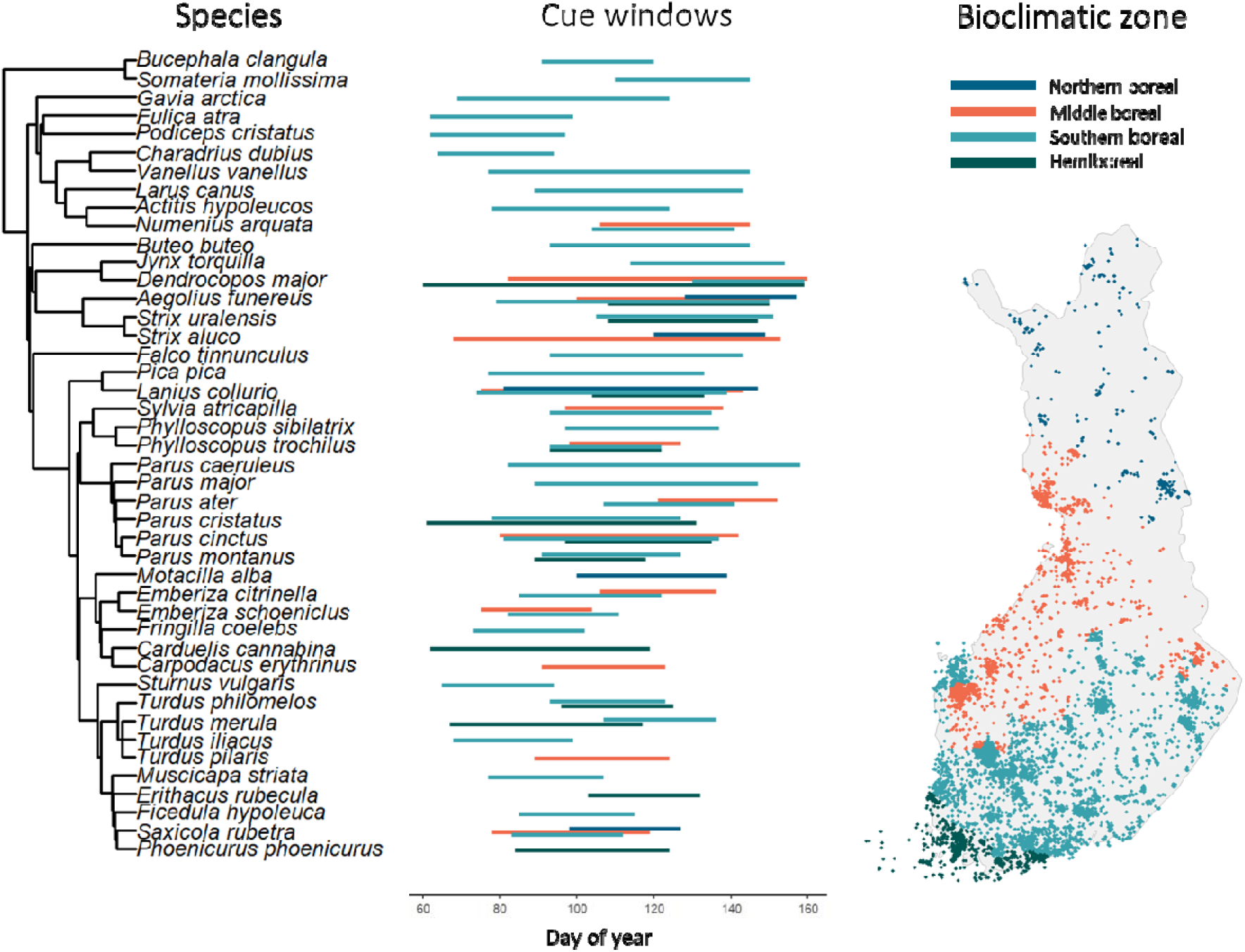
The best cue window for each of the 44 species that were retained for analyses and the spatial distribution of observations. Panel a) shows species ordered by phylogeny with the best cue window for each population, with populations colored by bioclimatic zone. Panel b) shows the distribution of species data on which the analyses were based with points colored by bioclimatic zone. See Fig. S1 for maps and histograms based on data at different stages of processing.

### 2.5. Population-level tests for shifts in reaction norms

For the retained populations we applied models to estimate (i) overall plasticity, and evidence for the (ii) *elevation adaptation hypothesis*, and (iii) *slope adaptation hypothesis*. To aid model convergence we first z-transformed the temperature variable (*t*) for each population. We then applied the within-subject centering approach (hereafter WSC; van de Pol & Wright 2009), wherein t_i j_ is the temperature for the population in year *i* and 5-year period *j*, *t^_j_* is the estimate of the mean temperature in a 5-year period and *t_i j_* - *t^_j_* is the within subject deviation in temperature experienced in year *i*. We anticipated that the slope β_W_ *t_i j_* - *t^_j_* will primarily capture plasticity (aim 1), whereas *β*_B_*t^_j_* was assumed to capture plasticity plus any contribution of elevation adaptation. Thus, where *β*_B_ departed from β_W_ this was assumed to estimate the adaptive evolution of the reaction norm elevation (aim 2), as in equation 1:

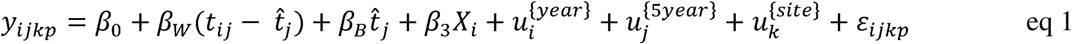

Here, y_tijkp_ represents the phenology (FELD) for year *i*, 5-year period *j*, site *k* and individual *p*. *β*_0_is the overall intercept, with *β*_3_ estimating the effect of year as a covariate (*X*) to detrend the data. □inline] and □inline] capture year, between 5-year subject, and site random effects, respectively, which are each estimated in the model. ∈_tijkp_ corresponds to the residual. All random effects assume a normal distribution with mean 0 and variance that is estimated in the model.

In the model represented in equation 1, where *β_W_* and *β*_B_ are both negative, then *β*_B_ < *β_W_* is consistent with co-gradient adaptation and *β*_B_ > *β_W_* is consistent with counter-gradient adaptation (Conover & Schultz, 1995, though this framing is more commonly applied in a spatial rather than purely temporal context) . To estimate the difference between the two slopes (*β*_B_- *β_W_*, i.e, our estimate of the contribution of elevation adaptation; aim 2; Fig. 1d-e) and its standard error, we can re-arrange equation 1 to obtain equation 2.

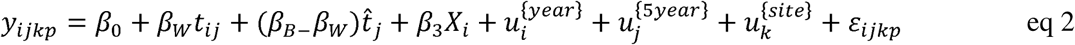

To test whether the reaction norm slope has changed over time (aim 3, *slope adaptation hypothesis*, Fig. 1f-g), we used a third model that included an interaction between year and the within-5-year period response:

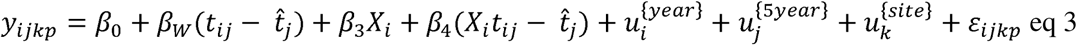

Assuming that *β*_w_ < 0, where the interaction (*β*_4_) is negative this would indicate that the reaction norm is becoming more steeply negative over time (Fig. 1g), whereas a positive value would indicate a population tending toward more positive (i.e., less steeply negative) reaction norm (Fig. 1f). The within-5-year period effect (*β_W_*) in this model can also be used to estimate average plasticity at the midpoint of the time series (aim 1).

Hereafter, we refer to the models based on the corresponding equation: M1 and M3 for responding to aim 1 about plasticity, M1 and M2 for responding to aim 2 on elevation adaptation, and M3 for responding to aim 3 on slope adaptation. For the subsequent meta-analyses and when presenting our results, we back-transformed all estimates, so they can be interpreted as change in *days/°C or days/year*.

### 2.6. Meta-analysis of responses across populations

To identify the average signal across a large pool of populations, whilst taking the uncertainty in estimates into account, we adopted a phylogenetic meta-analytic approach, using the *rma.mv* function in the *metafor* R package (Viechtbauer, 2010). We considered the following model estimates in the meta-analyses: (aim 1) β_W_ from M1 and M3, which we interpret as the average plasticity reaction norm; (aim 2) the difference in slopes β_B-_β_W_ from M2, testing the *elevation adaptation hypothesis*; and (aim 3) the within-5-year period by year interaction term in M3 (β_l_), testing the *slope adaptation hypothesis*. Additionally, we conducted a meta-analysis on the year effect (β_3_ from all three models; identical in M1 and M2) to assess whether we have evidence for a general temporal shift in phenology for reasons other than the temperature terms considered.

For each of the above effect sizes, we fitted a meta-analysis model including *Bioclimatic Zone* as a fixed effect (modifier) to estimate differences in effects sizes between the zones, *Phylogeny* as a random term to account for non-independence stemming from evolutionary relationships, and *Species* as a random effect to account for species-specific variation beyond phylogenetic relatedness. We used the back-transformed estimates as measures of effects size and their squared standard error as a measure of uncertainty in the effect size.

To test whether the effect of phylogeny was significant, we fitted a version of each meta-analytic model for aims 1-3 that did not include *Phylogeny* as a random effect and compared it, using AIC, to the main model. Additionally, we extracted the two estimated random effect variance components (σ²) from the main model. These variance components reflect the extent of (i) phylogenetic effects, and (ii) between-species heterogeneity that is not explained by the fixed effects nor accounted for by the phylogenetic random effect. We also estimated the phylogenetic signal (phylogenetic heritability) as the phylogenetic variance divided by the sum of the phylogenetic variance and the species variance (Lynch, 1991).

### 2.7. Testing the moderating effect of traits

To assess the effects of life-history traits on variation in adaptation of the reaction norm elevation or slope (aim 4), we included migratory strategy, average FELD per species, and generation length as moderators in corresponding meta-analyses models. A variance inflation factor test indicated low to negligible multicollinearity between the bioclimatic zones and the three trait variables (VIF < 4). We used AIC to compare the models including traits to the main meta-analysis models without traits.

### 2.8. Robustness test of potential shifts in cue timing

As our correlation-based inference approach is sensitive to the assumption that we have identified the causal cue for phenology (Tansey et al., 2017), we also examined whether there are shifts in FELD over time that are not captured by temperature effects (testing the effect of year, described in Section 2.7.), and whether the timing of the temperature cues’ themselves have shifted over time. This robustness test is described in Text S4.

All data processing and modelling was conducted in the R *environment* (R Core Team 2024, versions 4.1.1. and 4.4.1). Data and code are available in Hällfors et al. (2026).

## 3. Results

### 3.1. Cue windows

We found little evidence for consistent differences in the start or duration of the cue window between the bioclimate zones or across regions of the phylogeny (Fig. 2). Most windows were in the period from day 77–132 (i.e. 18 March – 12 May in non-leap years, based on 1^st^ and 3^rd^ quartile of start and end days, respectively) and the average duration was 44 days with a median of 39 days. We found no evidence consistent with a shift in the cue window timing (Text S4; Fig. S10, Table S1). After removing populations for which cue windows could not be reliably identified (Section 2.5.), we found that more reliable cue windows were detected more frequently in populations with more years of observations and with a longer time span of observations (Fig. S4).

Spring temperature in the breeding observation sites across days of the year 60–170 (the time window across which we identified thermal cue windows for the 77 species and 141 populations that these were initially estimated for) increased by 2°C (0.03 ± 0.0004°C /year) during the study period (Fig. S2), while within year variation in temperature during days of the year 60–170 (coefficient of variation; cv) decreased (−0.0005 cv units; Fig. S2). The increase in temperature was also similar in the observation sites in the different bioclimatic zones, with 2°C warming across the study period in the hemiboreal and southern boreal zones, 1.9°C in the northern boreal zones, and 1.8°C in the middle boreal zone. Over time, average temperature increased during the identified population-specific cue windows for the focal 69 populations (mean estimate = 0.04 C°/year, p < 0.001) with less warming in the southern and middle boreal zones compared to the hemiboreal (Fig. S3).

### 3.2. Aim 1: *Thermal plasticity*

The average estimated phenological plasticity across populations was -2.5 days °C□¹ (95% CI: -2.8, -2.1; Table 1, Fig. 3). The effects varied significantly by bioclimatic zone (ΔAIC = 8.4 when compared to a model without bioclimatic zones as moderators), with the strongest plasticity in the hemiboreal zone (−3 days °C□¹) and the weakest plasticity in the northern boreal zone (−2 days °C□¹). We found substantial phylogenetic variance in plasticity (σ^2^ =0.46) but no among-species variance, such that phylogenetic signal was strong (1.0). Phenological plasticity based on M3 were qualitatively similar (Table S2 and Fig. S7). For most populations (94%), the evidence for advanced phenology was robust, with confidence intervals excluding zero (Table S3). Examples of populations with especially strong phenological plasticity were *Carduelis cannabina* in SB (−9.5±2.5 days °C□¹; 95% CI: -14.4 to -4.6]), *Sylvia atricapilla* in SB (−6.1±1.5 days °C□¹; 95% CI: - 9.0 to -3.2), *Podiceps cristatus* in HB (−6±2 days °C□¹; 95% CI: -10.0 to -2.0), *Somateria mollissima* in HB (−5.9±1.8 days °C□¹; 95% CI: -9.4 to -2.4), *Gavia arctica* in SB (−5.7±1 days °C□¹; 95% CI: -7.6 to -3.7), and *Emberiza citronella* in SB (−5.1±1.3 days °C□¹; 95% CI: -7.7 to - 2.6).

**Figure 3.**
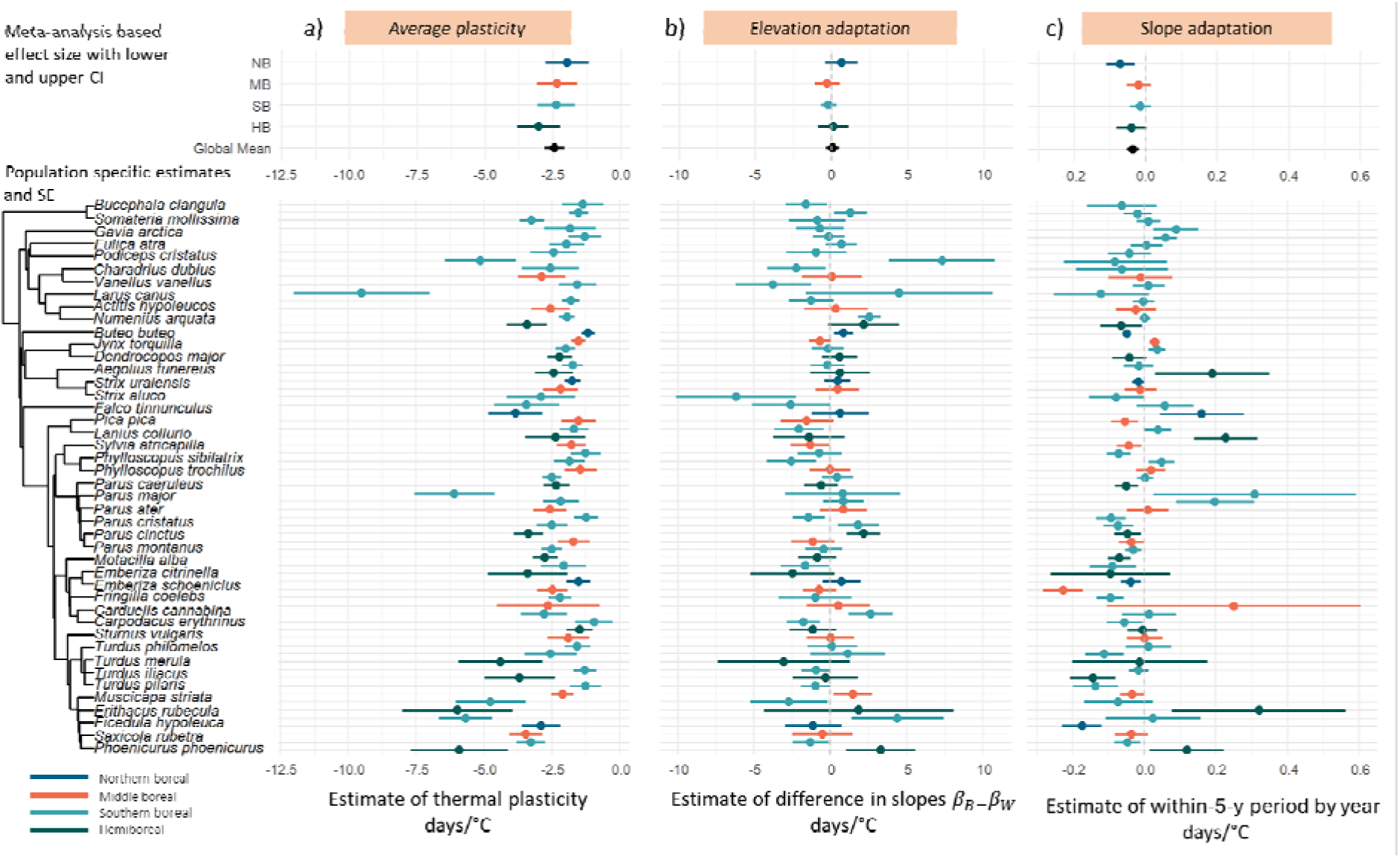
Test of (aim 1) thermal plasticity, (aim 2) *elevation adaptation,* and (aim 3) *slope adaptation*. Uppermost parts of panels show meta-analysis-based effects sizes with lower and upper confidence intervals per bioclimatic zone and the global mean of a) aim 1 on average plasticity (within-5-y period effect ( parameter from M1; within-5-y period effect ( from M3 presented in Fig. S7); b) aim 2 *on the elevation adaptation hypothesis* (the difference in slopes between versus within-5-y period ( ) from M2); and c) aim 3 on the *slope adaptation hypothesis* (the within-5-y period by year interaction term ( ) from M3). The lower parts of the panels show population-specific estimates and standard errors of respective aims. Boxplots and histogram of population-specific estimates are shown in Fig. S6.

**Table 1.**
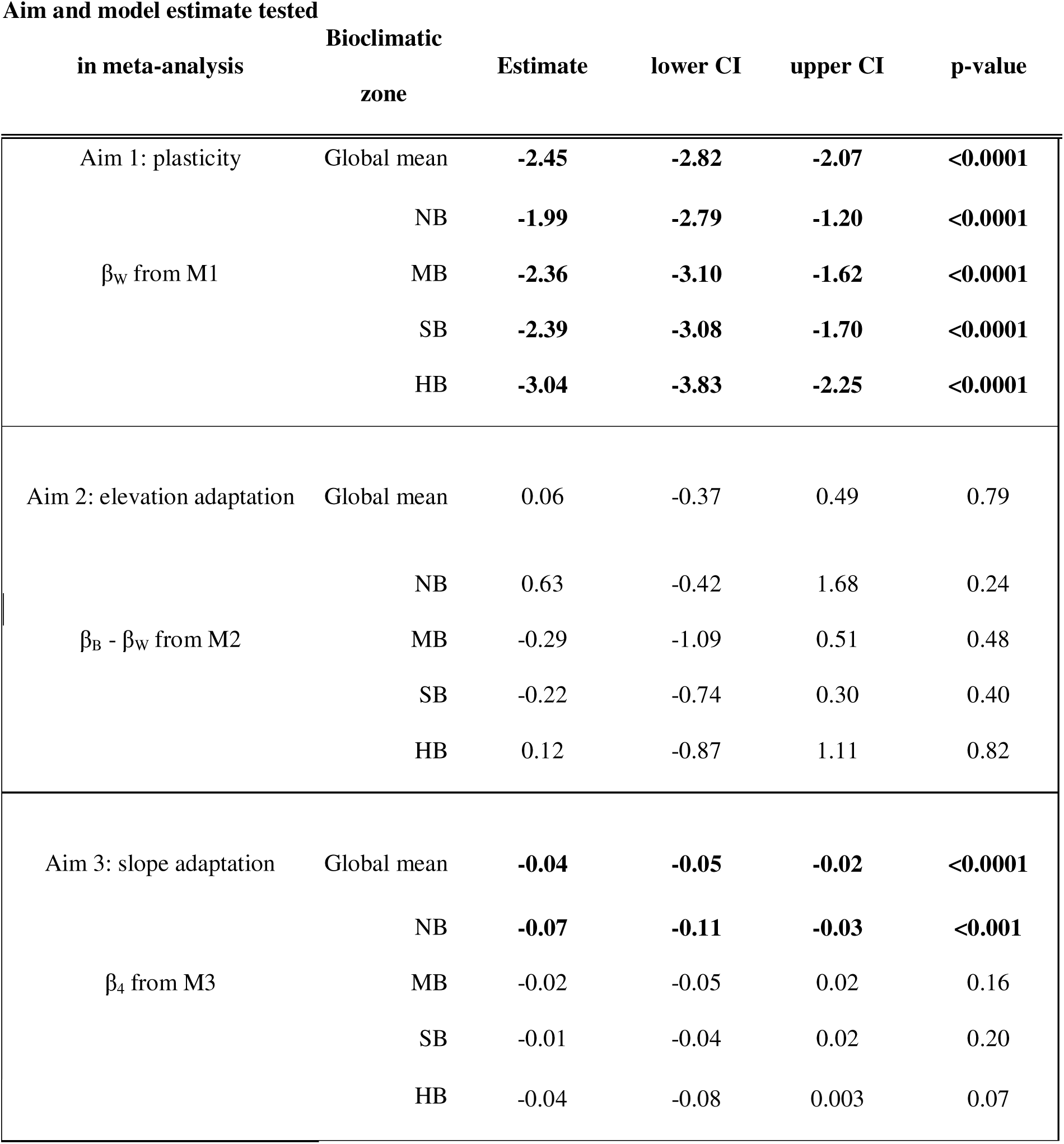
Summaries of meta-analysis models testing aim 1 on average plasticity (within-5-y period effect (β _W_) parameter from M1; within-5-y period effect (β _W_) from M3 presented in Table S2); aim 2 *on the elevation adaptation hypothesis* (the difference in slopes between versus within-5-y period (β _B-_β _W_) from M2); and aim 3 on the *slope adaptation hypothesis* (the within-5-y period by year interaction term (β’_4_) from M3). The results are visualised in Fig. 3. The column “Aim and model estimate tested in meta-analysis” indicates which aim (1-3) and model estimate was used a as response in the meta-analysis, and from which of the three models it was derived. The *BZ* column indicates which zone the estimate refers to, or whether it is the global mean estimate across zones. The estimates for each level of the bioclimatic zone have been adjusted to show the actual effects per level, not the difference to the baseline level. Statistically significant effects in bold. For aim 1, σ^2^ indicating the estimated variance of the random effects species and phylogeny were 0 and 0.49, respectively. The corresponding σ^2^ for aim 2 and 3, respectively, were 0.51 and 0, and 0 and 0. An AIC comparison to a model version excluding phylogeny indicated a negligible difference between the models with ΔAIC=0.76, -2, and -2 for aims 1, 2, and 3, respectively, with a negative AIC value indicating, in this case, a negligibly better fit for the model without a phylogeny. Funnel plots are shown in Fig. S5. NB= Northern boreal; MB= Middle boreal; SB= Southern boreal; HB= hemiboreal; CI= 95% confidence interval.

### 3.3. Aim 2: *Test of reaction norm elevation adaptation*

Across species, we found no evidence in support for *adaptation of reaction norm elevation* (Est. = 0.06 days °C□¹ , 95% CI: -0.37, 0.49). This means that, on average, the long-term reaction norms were not different to those over the short-term (as in Fig. 1c). There was also no significant difference between effects estimated for the bioclimatic zones (Fig. 3, Table 1). However, we found substantial among-species variance (σ^2^ = 0.51; Table 1), consistent with the true value of β_B_ - β_W_ being positive for some species and negative for others (Table S3). The phylogenetic signal was low (0), suggesting little resemblance in difference in short- vs. long-term slopes among closely related species. Only two populations showed clear evidence for a change in reaction norm elevation with confidence intervals excluding zero (counter-gradient response; 3%; Table S3). Several other species had positive or negative point estimates with reasonably precise estimates (small standard errors relative to effect size), although their confidence intervals did not always overlap zero. Examples of species with relatively strong co-gradient evolution of the reaction norm elevation (as in Fig. 1d) include *Erithacus rubecula* in SB (−6.2±3.9 days °C□¹; 95% CI: -13.9 to 1.5), *Fringilla coelebs* in SB (−3.7±2.5 days °C□¹; 95% CI: -8.6 to 1.1), and *Fulica atra* in SB (− 2.7±2.5 days °C□¹; 95% CI: -7.7 to 2.3), whereas species with signals of counter-gradient evolution of the reaction norm elevation (as in Fig. 1e) were *Emberiza citrinella* in SB (7.3±3.5 days °C□¹; 95% CI: 0.5 to 14.2), *Gavia arctica* in SB (4.4±3 days °C□¹; 95% CI: -1.5 to 10.3), *Somateria mollissima* in HB (3.3±2.3 days °C□¹; 95% CI: -1.2 to 7.8), and *Musciscapa triata* in SB (2.6±0.7 days °C□¹; 95% CI: 1.1 to 4.0).

### 3.4. Aim 3: Test of reaction norm slope adaptation

Across species, we found support for the *slope adaptation hypothesis* with plasticity increasing over the study period by -0.04 days °C□¹ (95% CI: -0.05, -0.02). With an average plasticity of -2.5 this global effect size of -0.04 means that, on average, the slope has become steeper by -2.6 days °C□¹ over the study period, being on average -2.5 days °C□¹ in the beginning of the study period and - 5.1 days °C□¹ in the end. The slope adaptation was especially evident in the northern boreal zone (−0.07 days °C□¹; Fig. 3c, Table 1) meaning that populations in the most northern parts of the study area had responded relatively more sensitively to temperature toward the present, by advancing their breeding timing more compared to the beginning of the 1960’s in equally warm years. Despite the average plasticity of populations in the northern boreal zone being weaker than across the whole study area (−2 days °C□¹ ), on average, plasticity in NB had become steeper by -4.5 days °C□¹ over the study period, being on average -2 days °C□¹ in the beginning of the study period but -6.5 days °C□¹ in the end. We found no support for among-species variance in slope adaptation nor any phylogeny-related variance (Table 1), but the phylogenetic signal was moderate (0.47), indicating partial clustering of year and short-term slope interactions among related species. Ten populations showed clear evidence for a steepening of reaction norm slope (14%), with confidence intervals excluding zero, while two showed clear evidence for evolving a shallower slope (3%; Table S3). Several other species had positive or negative point estimates with reasonably precise estimates (small standard errors relative to effect size), although their confidence intervals still overlapped zero. Examples of species with a strong steeping of reaction norms (as in Fig. 1f) were *Parus montanus* in MB (−0.23±0.06 days °C□¹; 95% CI: -0.3 to -0.1), *Bucephala clangula* in NB (−0.17±0.06 days °C□¹; □¹; 95% CI: -0.3 to -0.06), *Larus canus* in HB (−0.14±0.06 days °C□¹□¹; 95% CI: -0.3 to -0.02), and *Actitis hypoleucos* in SB (−0.14±0.06 days °C□¹ □¹; 95% CI: -0.3 to - 0.01), while examples of species with shallowing of reaction norms (as in Fig. 1g) include *Podiceps cristatus* in HB (0.32±0.24 days °C□¹□¹; 95% CI: -0.2 to 0.8), *Sylvia atricapilla* in SB (0.31±0.28 days °C□¹□¹; 95% CI: -0.2 to 0.9), *Turdus iliacus* in HB (0.22±0.09 days °C□¹□¹; 95% CI: 0.06 to 0.4), and *Ficedula hypoleuca* in MB (0.03±0.01 days °C□¹□¹; 95% CI: 0.0 to 0.06).

When we, as a robustness test, also fitted models with data based on more stringent selection criteria (Text S3), the results showed the same direction and a slightly stronger magnitude for aim 1, some more variation in magnitude for aim 2, and almost identical magnitude for aim 3 (Fig. S8; Table S4).

### 3.5. Aim 4: Effects of traits

To test the explanatory power of species traits on signals of genetic adaptation of the reaction norm, we compared the meta-analysis models for aim 2 and 3 to corresponding ones including three traits as moderators. Model comparison of the test of the *elevation adaptation hypothesis* (aim 2) the *slope adaptation hypothesis* (aim 3) showed that including traits did not improve model fit (ΔAIC = 2.95 and 3.97, in favour of the non-trait model for aims 2 and 3, respectively) This means that neither adaptation of elevation nor slope was explained by any of the tested species’ traits (Table 2, Fig. S9).

**Table 2.** Model statistics for meta-analysis responding to aim 4 on effect size of traits on change in a) elevation (the difference in slopes between versus within-5-y period β_B_ - β_W_ from M2) and b) slope (the within-5-y period by year interaction term (β_4_) from M3). In contrast to the non-trait model for aim 2 for which we found high phylogenetic variance but no among-species variance (Table 1), we found no phylogenetic structure (σ^2^ = 0) but high among-species variance (σ^2^=0.43) in the trait model for elevation adaptation. Like in the non-trait model for aim 3, we found no species nor phylogeny related differences, with a ^2^ of 0 for both among-species variance and phylogeny in the trait model for slope adaptation. Resident and the hemiboreal zone are captured by the intercept. SB = southern boreal, MB = middle boreal, NB = northern boreal. See plotted model predictions in Fig. S9.

| Parameter | Estimate | lower<br>CI | upper<br>CI | z-<br>value | p-value |
| --- | --- | --- | --- | --- | --- |
| Intercept | 3.38 | -2.87 | 9.64 | 1.06 | 0.29 |
| Zone SB | -0.38 | -1.45 | 0.69 | -0.70 | 0.49 |
| Zone MB | -0.37 | -1.58 | 0.83 | -0.61 | 0.55 |
| Zone NB | 0.41 | -0.97 | 1.78 | 0.58 | 0.57 |
| Migration type - short | -0.38 | -1.64 | 0.87 | -0.60 | 0.56 |
| Migration type - long | 0.62 | -0.94 | 2.18 | 0.78 | 0.44 |
| Generation length | -0.11 | -0.37 | 0.15 | -0.80 | 0.43 |
| Average day of year | -0.02 | -0.06 | 0.02 | -0.92 | 0.36 |

b) Zone and trait effects on slope adaptation (Aim 3)
| Parameter | Estimate | lower<br>CI | upper<br>CI | z-<br>value | p-value |
| --- | --- | --- | --- | --- | --- |
| Intercept | 0.06 | -0.19 | 0.31 | 0.47 | 0.64 |
| Zone SB | 0.03 | -0.01 | 0.06 | 1.44 | 0.15 |
| Zone MB | 0.02 | -0.02 | 0.06 | 1.00 | 0.32 |
| Zone NB | -0.03 | -0.08 | 0.01 | -1.61 | 0.11 |
| Migration type - short | 0.01 | -0.04 | 0.06 | 0.31 | 0.76 |
| Migration type - long | 0.04 | -0.02 | 0.10 | 1.44 | 0.15 |
| Generation length | 0.00 | -0.01 | 0.01 | 0.10 | 0.92 |
| Average day of year | 0.00 | 0.00 | 0.00 | -1.00 | 0.32 |

## 4. Discussion

In this study, we have demonstrated how long-term observational data can be used to gauge signals of evolutionary adaptation in plasticity over time. We found exciting evidence for evolution of reaction-norm slope for several bird species, with phenological sensitivity to temperature on average steepening over time. This change was particularly evident in the northern boreal zone, where the plastic slope was estimated to have steepened 4.5 days °C□¹ between the beginning and end of the time series. We also found clear evidence that breeding phenology of birds is thermally plastic in general, with phenology advancing by 2.5 days□°C□¹ on average with the realized change in thermal conditions over the study period, with evidence for among species phylogenetic variation in the slope. We detected no meta-analytic support for adaptation of the reaction-norm elevation, though we did find among species variation in slope, such that the reaction norm elevations of some species may be adapting in both co- and counter-gradient directions (Fig. 1d-e).

### Aim 1: Average phenological plasticity

We found differences in average plasticity between the bioclimatic zones, with higher plasticity in the south (−3.04 days□°C□¹ in HB) which gradually decreased towards the north (−1.99 days□°C□¹ in NB; Table 1). This difference may be due to restrictions towards plastic advances further to the north caused by, e.g., arrival time of migrants, a relatively stronger dependency on day length cues, and variability in snow melt timing cues. The average temperature sensitivity (−2.5 days□°C□¹) and the compelling evidence from across the studied populations implies that the vast majority of populations have plastically advanced their egg-laying timing by approximately 5 days over the study period, given the ∼2°C increase in temperature during spring. Our zone-specific estimates showed spatial heterogeneity in realized plastic advances when combined with zone-specific realized thermal increases, with an average plastic advance of 5 days in the hemiboreal, 6.5 days in the southern boreal, 4.3 days in the middle boreal, and 3.8 days in the northern boreal. Overall, these plastic advances were consistent with the expectation that phenotypic plasticity on its own allows rapid tracking of a temperature-dependent optimum (Charmantier & Gienapp, 2014; Chevin et al., 2010; Merilä & Hendry, 2014), and the magnitude of plasticity was largely in line with previous studies on boreal bird breeding timing in response to climate change (Hällfors et al., 2020; Kluen et al., 2017)

### Aim 2: Elevation adaptation

We found no evidence that, across species, the average reaction norm elevation in short-term periods would differ from those across the long-term. This suggests no consistent, across-population adjustment of mean breeding timing beyond what plasticity explains. However, we found large variability between species for adaptation of the reaction norm elevation, although no individual populations showed robust signals for co-gradient evolution of the reaction norm elevation and only 3% of populations showed that for counter-gradient evolution. Although plasticity on its own may indeed be enough to help species adjust without genetic adaptation, two other factors could contribute to the lack of an overall signal in adaptation of the reaction norm elevation. If plastic responses buffer phenological lags relative to a moving optimum (Chevin et al., 2010; Tansey et al., 2017), this reduces the necessity for and contribution of genetic adaptation. To better understand the extent to which plasticity is adequate we need more estimates of how the optimum FELD is changing with spring temperature (e.g., Chevin et al., 2015; van Asch et al., 2013). The second factor that could explain the lack of signal is that a large part of the time series used for in this study (from 1961–2025) covers decades with weaker warming and hence weaker selection pressure, as the majority of climate warming, and thus higher potential selection pressure, has mainly occurred during the past three decades (Mikkonen et al., 2015). Repeating this approach with additional data or in the future as more observations from periods that have experienced strong climate change has accumulated, could increase our power to detect elevation adaptation.

The high among-species variance in reaction norm elevation shifts that we detected is consistent with idiosyncratic species-level signals, with some species shifting the elevation of the reaction norm, but we found strong evidence for this for only two individual populations. However, the absence of phylogenetic effects implies that these differences among species are not constrained by relatedness, and instead there may be other factors making evolutionary shifts in mean breeding timing more likely for some species. Indeed, several species showed co-gradient-type adaptation of the reaction norm, with short-term slopes being shallower than the long-term slope (Fig. 1d), while others showed indications of a counter-gradient-type adaptation over time, with steeper short-term slopes compared to the long-term slope (Fig. 1e). A next step to better understand these trends is to examine whether in warm years we see directional selection for earlier laying in the populations exhibiting co-gradient patterns and a tendency for selection to favour later laying birds in the counter-gradient cases. While we conducted a number of data selection steps and tests to avoid including populations for which cue identification was less certain, it is possible that some species use other cues than temperature (such as prey or seed abundance; Kanerva et al., 2020; Lehikoinen et al., 2011) or a combination of environmental cues for timing their breeding (Charmantier & Gienapp, 2014; Shutt et al., 2019). In addition to direct daily temperature, species may use photoperiod and other signs of an advancing spring, such as vegetation green up or thermal sums, to time their breeding initiation. If the timing of these do not change in relation to the timing of resources, or in case of higher variability in phenological optima, we would expect no selection of either the elevation nor slope of the reaction norm (Ramakers et al., 2019). Especially for migrating birds, identifying the correct cues in the breeding area can be difficult, although temperature is typically spatially autocorrelated over a large geographical area (Halkka et al., 2011).

### Aim 3: Slope adaptation

We found meta-analytical support for bird breeding timing having become more responsive to thermal conditions over time, with an overall change in thermal sensitivity of -2.6 days °C□¹ over the study period. In addition, 14% of population-specific estimates showed strong evidence of such a response. Specifically, the revealed negative interaction between year and temperature during short-term periods indicates a steepening of thermal reaction norms toward the present and is consistent with a scenario of adaptation of phenological plasticity, where populations experience selection because their plastic slope is lagging behind the optimum sensitivity and have therefore evolved steeper slopes over time.

We also found some evidence that plasticity of certain populations (3%) has evolved to become shallower. Previous studies on phenological evolution on winter moth (*Operophtera brumata;* van Asch et al., 2013), found some contribution through a shallowing of the reaction norm slope while a larger contribution stemmed adaptation of the elevation, together leading to moths hatching later in all temperatures, but relatively even later in warm temperatures to match the timing of bud burst in oak (*Quercus robur*), on which they feed. There is some evidence that Arctic plant and insect species that have been exposed to large temperature increases may be reaching limits to their plasticity (e.g., Gerlich et al., 2025; Iler et al., 2013). The low signals of among-species and phylogenetic variance that we found suggest that the population effect sizes that are consistent with a shallowing slope may be attributable to sampling variance.

Our study is, to our knowledge, the first to find extensive evidence for evolution of plasticity for multiple species and offers exciting opportunities for further in-depth studies on, e.g., those with the strongest evidence of such a response, such as *Parus major* in the hemiboreal zone in Finland. A study on the great tit (*Parus major*), found potential for evolution of the reaction norm elevation but not of its slope (Ramakers et al., 2019), with breeding timing expected to occur earlier over time but with no change in breeding slope. This was supported by evidence for a shift towards earlier optimal laying dates across the thermal range and additive genetic variation in reaction norm elevation, while no such evidence was found for slope. Interestingly, our results for *Parus major* in the hemiboreal zone are the opposite to what Ramakers et al. (2019) predict for the populations they studied in the Netherlands, as we, in addition to finding signals of increased plasticity, observed no evidence for evolution of the reaction norm elevation for this population. This difference may be due to different data types, time spans, and methodological approaches, but also to genetic differences between Dutch and southern Finnish populations of *Parus major* and their environmental contexts, with the latter alternatives being more likely based on us not finding evidence adaptation of the reaction norm slope for *Parus major* in the southern and middle boreal zones.

A steepening of the plastic slope is predicted under scenarios where there is an increase in the correlation between the environment of development (i.e. the cue used to determine plasticity) and the environment of selection (the environment that determines the optimum ; Gavrilets & Scheiner, 1993). Interestingly, this increasing plastic sensitivity to temperature was most evident in the northern boreal zone, where species overall showed the weakest plasticity. Indeed, insufficient plasticity may increase selection pressure, while at the same time not masking the genotype that selection can act upon (Charmantier et al., 2008; Oostra et al., 2018). However, strong phenotypic plasticity can also aid adaptive evolution in novel conditions, as it allows species to respond adaptively in the short-term and gives more time for evolution to act in the long term (Price et al. 2003; Sgró et al 2016). This, in turn, could explain why we found close to significant effects in the hemiboreal zone, where plasticity was highest. In the future, it would be important to explicitly study how levels of plasticity relate to signatures of reaction norm evolution, and whether more plastic populations tend to adapt their reaction norm elevations and slopes more rapidly, or vice versa. How such trade-offs will pan out in the future will be important to monitor, and the approach that we present in this paper offers a promising avenue for monitoring such changes using available and constantly accumulation observational data on fitness related traits such as species phenology.

### Aim 4: Trait effects

We expected to find that resident and short-distance migrants, early breeding species, and species with shorter generation times would show stronger signals of evolutionary adaptation in their breeding phenology. However, we found no evidence for such connections. Previous studies have shown that resident species and those that tend to breed earlier during the season advance their phenology more readily than others (Hällfors et al., 2020; Kluen et al., 2017). In the light of our results, this may be driven by plasticity alone with no recent adaptation specifically contributing to these differences between residents versus migrants and early versus late breeders. The fact that we found evidence for birds with shorter generation times to be more capable of evolving rapidly may be due to the low number of tested species, but also to that short- and long-lived birds may in fact not differ that much in their evolutionary potential. Also relatively long-lived birds have been found to show rapid evolutionary responses (Grant & Grant, 2002; Moiron et al., 2024) and this may be due to higher additive genetic variation in plasticity among longer lived species, which underpins evolution of plasticity, whereas short-lived birds may overall, like the early breeding species, be more plastic to begin with.

### Methodological Considerations and Limitations

We performed a series of robustness checks and sensitivity analyses to evaluate the reliability of cue window identification, the stability of thermal reaction norm estimates, and potential sources of statistical bias. These included filtering populations with uncertain cue windows, permutation tests, alternative quantile thresholds for cue identification, and two-period sensitivity analyses to test for temporal shifts in cue timing. Collectively, these analyses indicate that the identified thermal cue windows are generally robust, unlikely to have shifted substantially over time, and not driven by changes in temperature variability or sampling structure. At the same time, our approach entails important methodological considerations, including statistical coupling between within- and between-population effects (Phillimore et al., 2010; Westneat et al., 2020), sensitivity of slope estimates to sliding window selection, and biases to effect sizes introduced by retaining only populations with strong temperature signals. While these caveats do not appear to undermine the main conclusions, they are important for interpretation and for future applications of this framework. A detailed discussion of these methodological considerations and limitations is provided in Text S5.

## Conclusions

The results from our study show strong thermal plasticity in avian breeding phenology and evidence of emerging adaptation of plasticity, which we suggest is a consequence of shifting selection caused by climate change. To predict changes in biodiversity under ongoing and increasing climate change, understanding the role of plasticity versus genetic adaptation is critical. Our ability to do so is, however, halted by the limited knowledge on the extent to which different species rely on both or either mechanism (Charmantier & Gienapp, 2014), ultimately undermining our capacity to predict how communities and ecosystems will change over time. The findings of this study reinforce the importance of plasticity–adaptation interactions in understanding species responses to climate change and demonstrate how long-term observational data can help reveal potential cases of evolutionary adaptation that can be relevant when forecasting ecological responses in key life-history traits under continued environmental change. Such a macroecological assessment can, in addition to gauging signs of evolutionary responses across multiple taxa, allow identifying species for more in-depth experiments and causal testing. While observational studies based on individual and longitudinal data and experimental approaches will continue to play an essential role in robustly demonstrating relative contributions of plasticity versus evolutionary adaptation under climate change (Charmantier & Gienapp, 2014), our approach offers a gateway to monitor evolutionary signals in reaction norms using long-term observational data and even citizen-science-based observations on multiple species.

## Acknowledgements

We are grateful to the thousands of bird nest recorders that have reported their observations through the Finnish nest card scheme. The Research Council of Finland funded work of MHH (grants 330739 and 360742) and AL (grant 362647). MHH also acknowledges funding through the Finnish Strategic Research Council’s projects IBC-Carbon (312559) and MUST (358367), the Finnish Ministry of the Environment (the SUMI project, grant number VN/33334/2021; and the FEO project, grant number VN/12351/2021), and funding by the Jane and Aatos Erkko Foundation through the Research Centre for Ecological Change. We thank Andrea Santangeli, Bess Hardwick, and Tanja Lindholm at the Research Centre for Ecological Change for trait compilation, and Juha Aalto and Pentti Pirinen at the Finnish Meteorological Institute for help with accessing weather data. We acknowledge the use of CoPilot and ChatGPT for debugging and developing pieces of R code, for shortening and structuring parts of the abstract and discussion, and for screening the manuscript for inconsistent cross-referencing. The authors reviewed and edited the AI-generated content before including any of the suggestions in the work.

## Author contributions

MHH and AP conceived of the idea. AL provided data and interpretation of data and first egg-laying date estimations. MHH processed data in close collaboration with AL. MHH conducted statistical analyses in close collaboration with AP. MHH led the writing of the manuscript and AP and AL contributed essentially. All authors gave final approval of submitting the manuscript.

## Data availability statement

Data and code are available for peer review in Dryad (DOI:10.5061/dryad.t1g1jwtj6) and will be published upon acceptance of the manuscript.

