## Supplemental Data 1 for "Widespread evidence for plasticity and recent evolution of plasticity in the breeding phenology of Finnish birds"

Contents:

Figures S1-S10

Tables S1-S4

Texts S1-S5

References

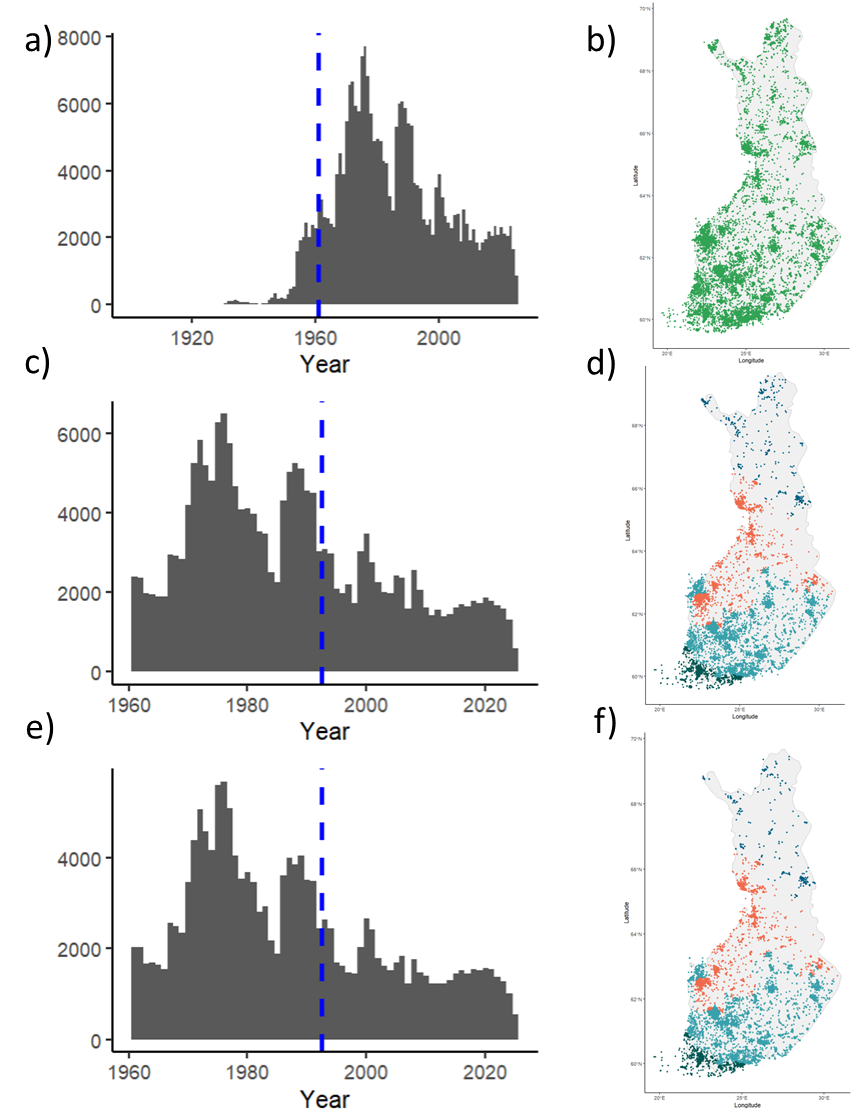

**Figure S1.** Original data and subsets of data for analyses. Panel a) is a histogram of the complete raw data set with bar height indicating annual number of monitored nests across Finland. For subsequent analyses we only included data from 1961 onwards (indicated by vertical blue line in panel a)), as the observation numbers at this point reach an overall higher level and since daily weather data were available from 1961 onwards. Panel b) is a map of Finland showing the spatial distribution of the complete nest card data set starting in 1901, including 249 807 recorded nests of 236 species out of all 250 breeding species in Finland. We further delimited these data as part of FELD estimation and removal of records prior to 1961, and included only species in bioclimatic zones with sufficient numbers of observations (see main text; *2.1.*). Panel c) is a histogram of the delimited observations and panel d) is a map of Finland showing the spatial distribution of the remaining data after, containing 190 338 observations of 141 populations of 77 species. These data were further delimited for the final analyses based on robustness of identified cue windows (see main text; *2.4.* and Text S3) leaving us with 69 populations of 44 species to be analyzed (data available in Hällfors et al. (2026)). Panel e) is a histogram of the analysed data and panel f) a map of the spatial distribution of the analysed data. The vertical blue line in panels c) and e) are placed at 1992.5 as observations from 1961-1992 were assigned into T1 and 1993-2025 to T2 when dividing the study period into two halves for the sensitivity analyses testing potential cues shifts over time (see main text; *2.8.* and Text S4). Colours in panel d) and f) indicate in which bioclimatic zone the observation was situated (teal= hemiboreal; light blue= southern boreal; coral= middle boreal; dark blue= northern boreal).

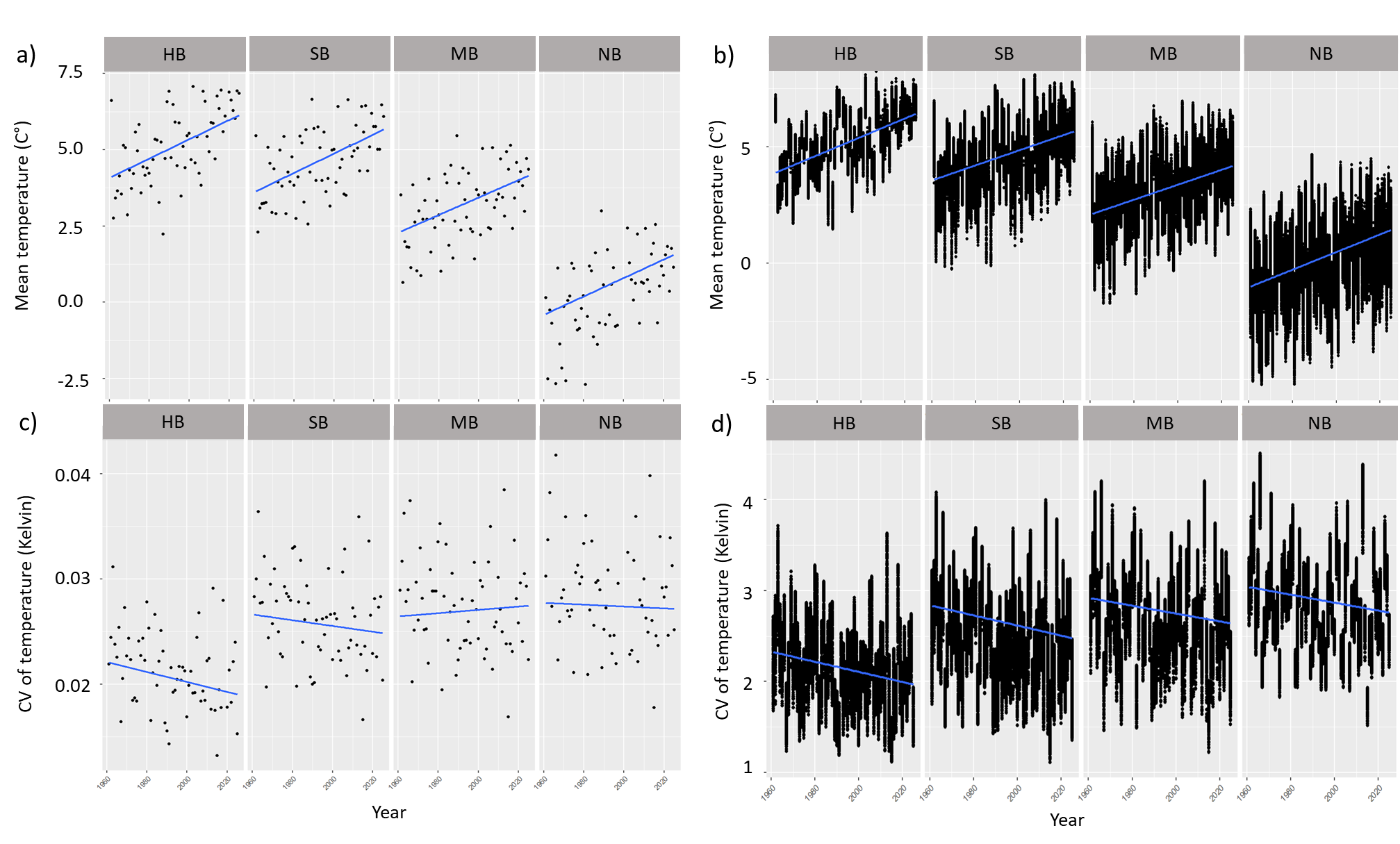

**Figure S2.** Change in temperature mean and variation over time. **Panel a)** shows mean temperature (°C) change over time in observation sites) for the 77 species and 141 populations that these were initially estimated for (Main text; section 2.1; Fig S1c-d), across days of year 60-170, the seasonal period during which we identified cue windows, separated in sub-panels by bioclimatic zone. We conducted linear regressions testing the effect of year on mean temperature during day of years 60-170, and they all showed statistical significance levels of <0.0001. **Panel b)** shows the same as panel a) but across all potential spatial sites in Finland (combinations of latitude and longitude). Temperatures in the observation sites a) across the whole of Finland have increased by 2°C during the study period (0.03±0.0004°C/year) while the b) overall change in temperature was 2.1°C (0.035±0.0002°C /year). This implies that the observations were, overall, done in sites that have warmed similarly to the average warming in the whole country. When examining models including an interaction term between year and bioclimatic zones there were significant differences only between the middle boreal and hemiboreal in thermal increase in the sampling sites (p<0.05), but a significant difference (p<0.05) between all zones compared to the reference zone HB for all potential sites in Finland. Separate models for each zone (data split per zone and only year as fixed effects) showed the following increases in temperature: the hemiboreal zone (HB) showed an increase of 2°C (0.03±0.002°C /year) in observation sites and 2.5°C (0.039±0.0004°C /year) in the whole HB zone (i.e., less warming in observation sites); in the south boreal zone (SB) we saw an increase of 2°C (0.03±0.0003°C/year) in the observation sites and 2.2°C (0.03±0.0002°C /year) in all potential spatial sites (i.e., slightly less warming in observation sites compared to across the whole zone); in the middle boreal zone (MB) we saw an increase of 1.8°C (0.03±0.0006°C/year) in the sampling sites and a 2.1°C (0.03±0.0002°C /year) increase in all potential sites (slightly stronger warming trend in the observation sites as to in the all potential sites); and in the north boreal zone (NB) we saw a thermal increase of 1.9°C (0.03±0.001°C/year) in the observation sites and a 2.4°C (0.04±0.0003°C /year) increase in all potential sites (i.e., less warming in the observation sites). Thus, we can conclude that there is no systematic bias across all parts of the country regarding where the observations of nests were done vis-à-vis the overall warming trends, but there were overall differences between all potential and sampling sites that may mirror preferences of the nest visitors towards sites that are warming less rapidly than on average, especially in the NB zone. **Panel c)** shows change over time in the coefficient of variation (CV) of temperature (Kelvin) in the observation sites across days of year 60-170. **Panel d)** shows the same as a but across all potential spatial sites. All linear regressions testing the effect of year on the coefficient of variation during day of years 60-170 showed statistical significance levels of <0.0001 if not mentioned otherwise. Temperature variation in the observation sites (c) had decreased by −0.0005 CV units during the study period (−7.6×10⁻^6^±1.3×10⁻⁶ CV units/year) while the overall change in variation in all potential spatial sites was -0.31 CV units (−4.8×10⁻^3^±5.3×10⁻^5^ CV units /year). This implies that the observations were, overall, done in sites that have stayed more similar in variation over spring compared to the average change in variation in the whole country. When examining models including an interaction term between year and bioclimatic zones showed significant difference between the sampling sites in the southern, middle and northern boreal from the intercept level hemiboreal, and for all potential sites in the middle and northern boreal compared to the hemiboreal. While the overall trend of decreased variation across spring was mirrored in zone-specific changes (modelled separately for each zone with only year as a fixed effect), the exception was the middle boreal zone, where the variation had increased in the observation sites while it had decreased across all other observation and potential spatial sites. Change in CV over the study period in HB was -0.003 CV units for observation sites (-4.7×10⁻⁵±4.5×10⁻⁶ CV units/year) and 0.005 for all potential sites (-0.006±0.0002 CV units/year); in SB it was -0.002 CV units (-2.7×10⁻^5^±1.5×10⁻⁶ CV units year) for observation sites and -0.005 CV units (-5.6×10⁻^3^±8.2×10⁻^5^ CV units/year) for all potential sites; in the MB the variation increased by 0.001 CV units (1.6×10⁻⁵±2.6×10⁻⁶ CV units/year) in the observation sites but decreased by -0.004 CV units (-4.3×10⁻^3^±8.9×10⁻^5^ CV units/year) in all potential spatial sites; finally in the NB the CV did not change in sampling sites (-8.8×10⁻^6^±7.4×10⁻⁶ CV units/year; p=0.23) but changed by -0.004 in all potential spatial sites (-4.4×10⁻^3^±9.6×10⁻^5^ CV units/year). This implies that, overall, birds may have come to experience more similar conditions in springtime, which can make changes in plasticity more difficult to detect. However, as shown by the difference between change in variation in the sampling sites versus overall, the nest visits in the MB and NB zones may be biased to areas in which variation over spring has remained more similar over time.

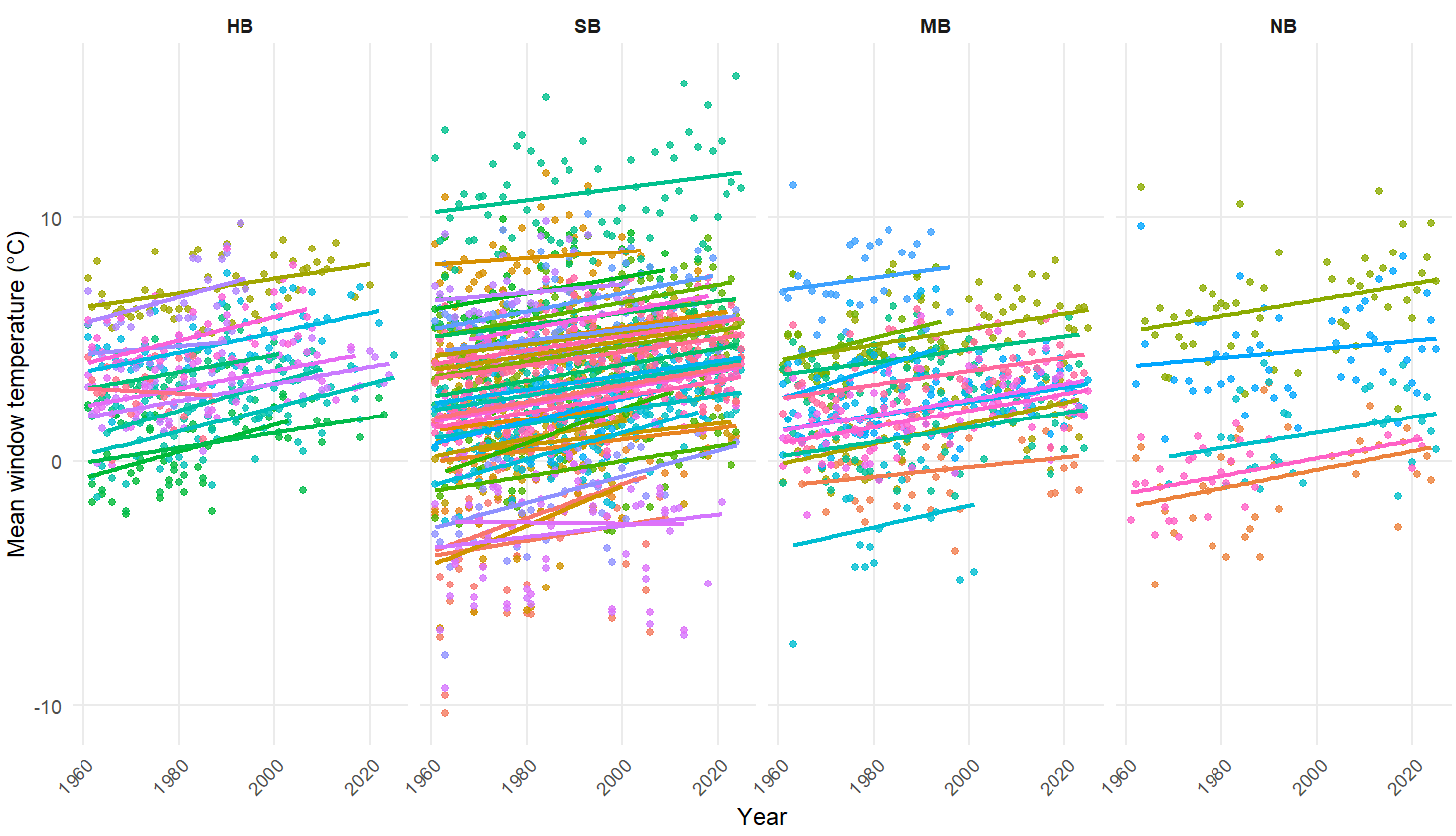

**Figure S3.** Temperature change over time during the best cue window for each of the 69 populations of 44 species (different colors) analyzed in the study. Bioclimatic zones separated by panels; HB= hemiboreal zone; SB= southern boreal; MB= middle boreal; NB= northern boreal. Temperature during the population-specific cue windows had increased over time (Est.= 0.04 °C/year, p<0.001) with less increase in the southern (estimated difference in slope compared to the intercept level=-0.01+-0.005, p<0.05) and middle boreal zones (estimated difference in slope compared to the intercept level =-0.02+-0.006, p<0.01) compared to the hemiboreal (intercept level), while the difference in estimated slope did not differ between the northern boreal and hemiboreal zones (estimated difference in slope=0.02+-0.008; p=0.77).

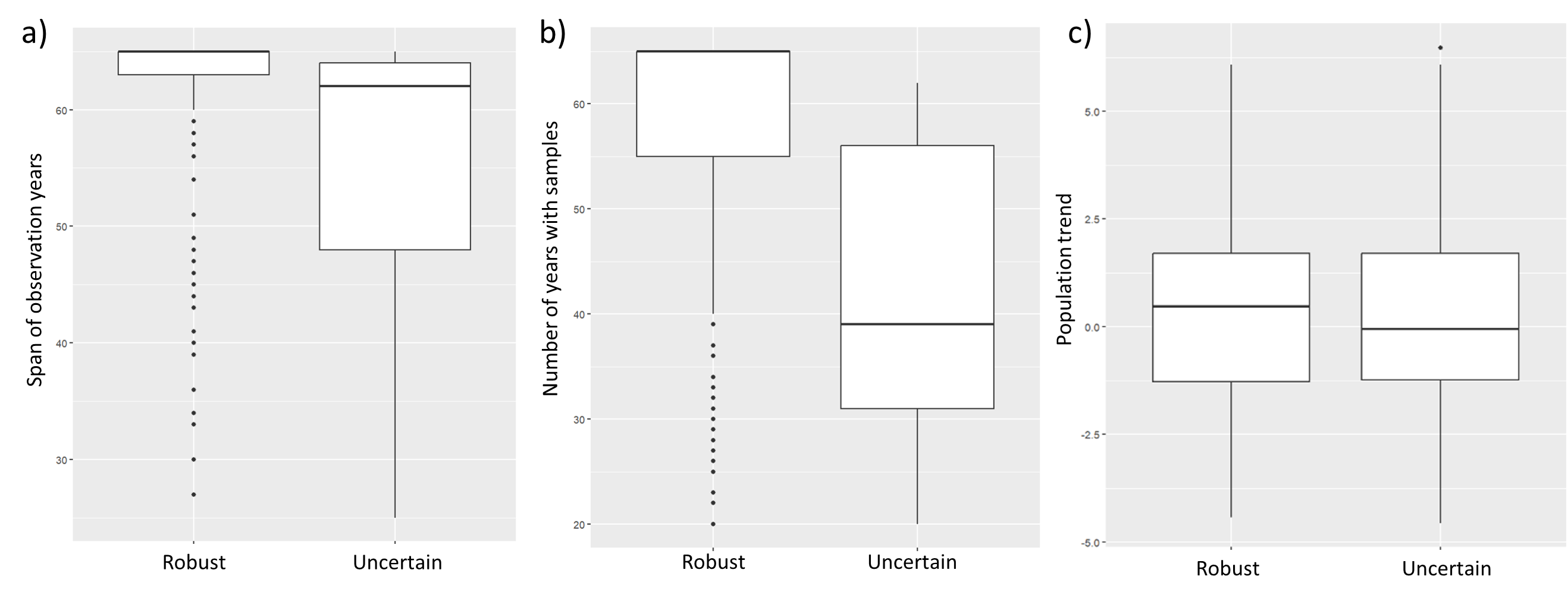

**Figure S4.** Boxplot showing differences between populations for which we found relative certainty in the identified window (robust; 69 populations of 44 species) versus those for which other cue windows or random cue windows explained the data equally well or better (uncertain; 72 populations of 33 species) compared to the a) span of years for which we had observations and the number of years with observations, b) number of years for which we had samples, and c) population trends (NB! Available only for 57/77 species; 109/141 populations). The difference between the groups was statistically significant for year span and the number of sampled years, indicating that populations with a shorter span of observations across the study period (Est. -6.1 years; p<0.001) and less sampled years (Est. -17.1 years, p<0.001) tended to have uncertain cue windows. There was no difference between in species’ population trend between populations with robust and uncertain cue windows (Est. -0.07, p=0.87).

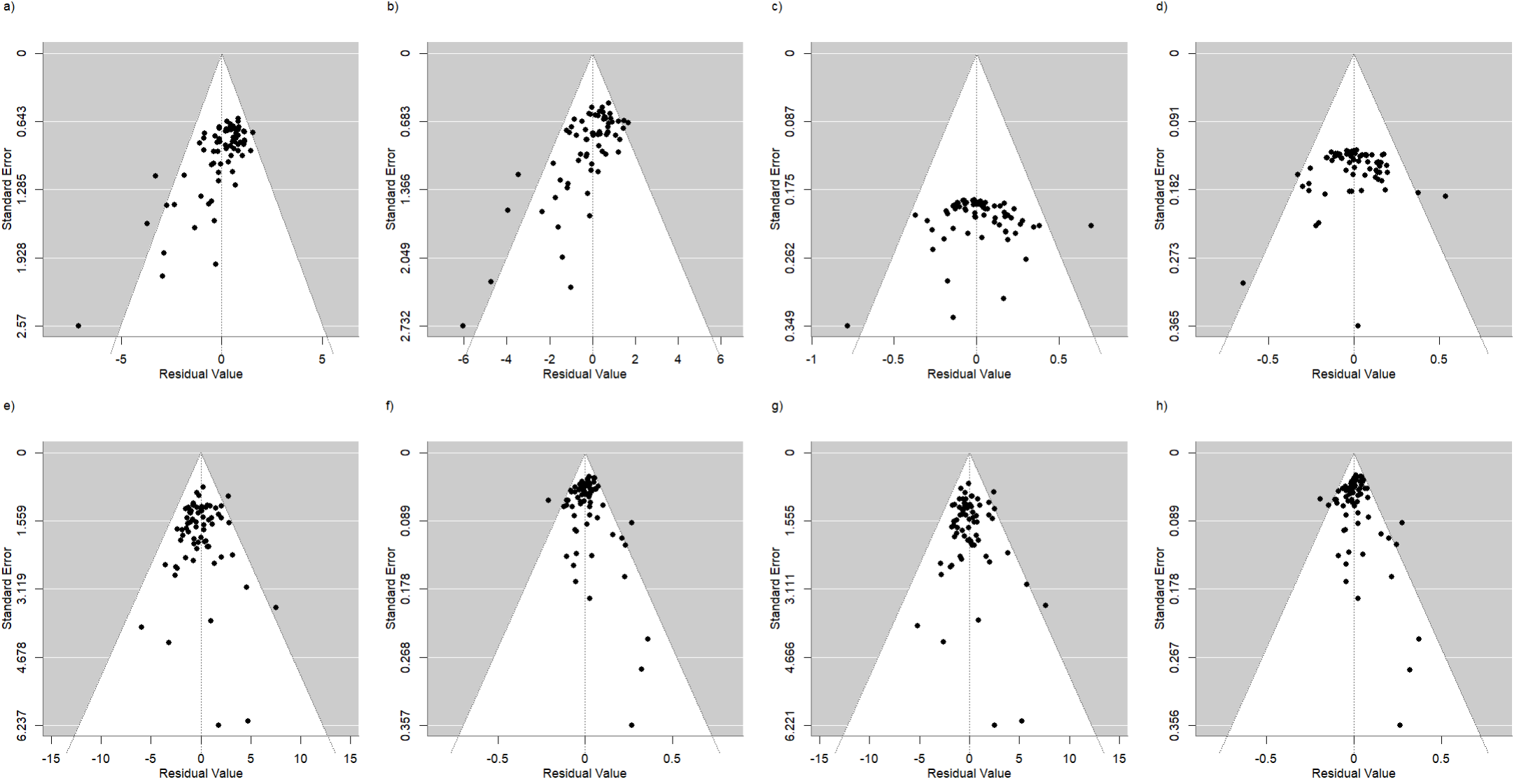

**Figure S5.** Funnel plots based on meta-analyses with the standard error on the y axes and residual error on the x-axes. Panels show funnel plots on meta-analysis models on a) average plasticity (within-5-y period effect $\beta_{W}$ parameter from M1), b) average plasticity (within-5-y period effect $\beta_{W}$ parameter from M3), c) year effect ($\beta_{3}$ from M1 and M2 (identical)), d) year effect ( $\beta_{3}$ from M3), e) difference in reaction norm elevation between versus within-5-y period ($\beta_{B-}\beta_{W}$ from M2), f) change in reaction norm slope with time (the within-5-y period by year interaction term $\beta_{4}$ from M3), g) difference in reaction norm elevation between versus within-5-y period in the trait model, and h) change in reaction norm slope with time in the trait model.

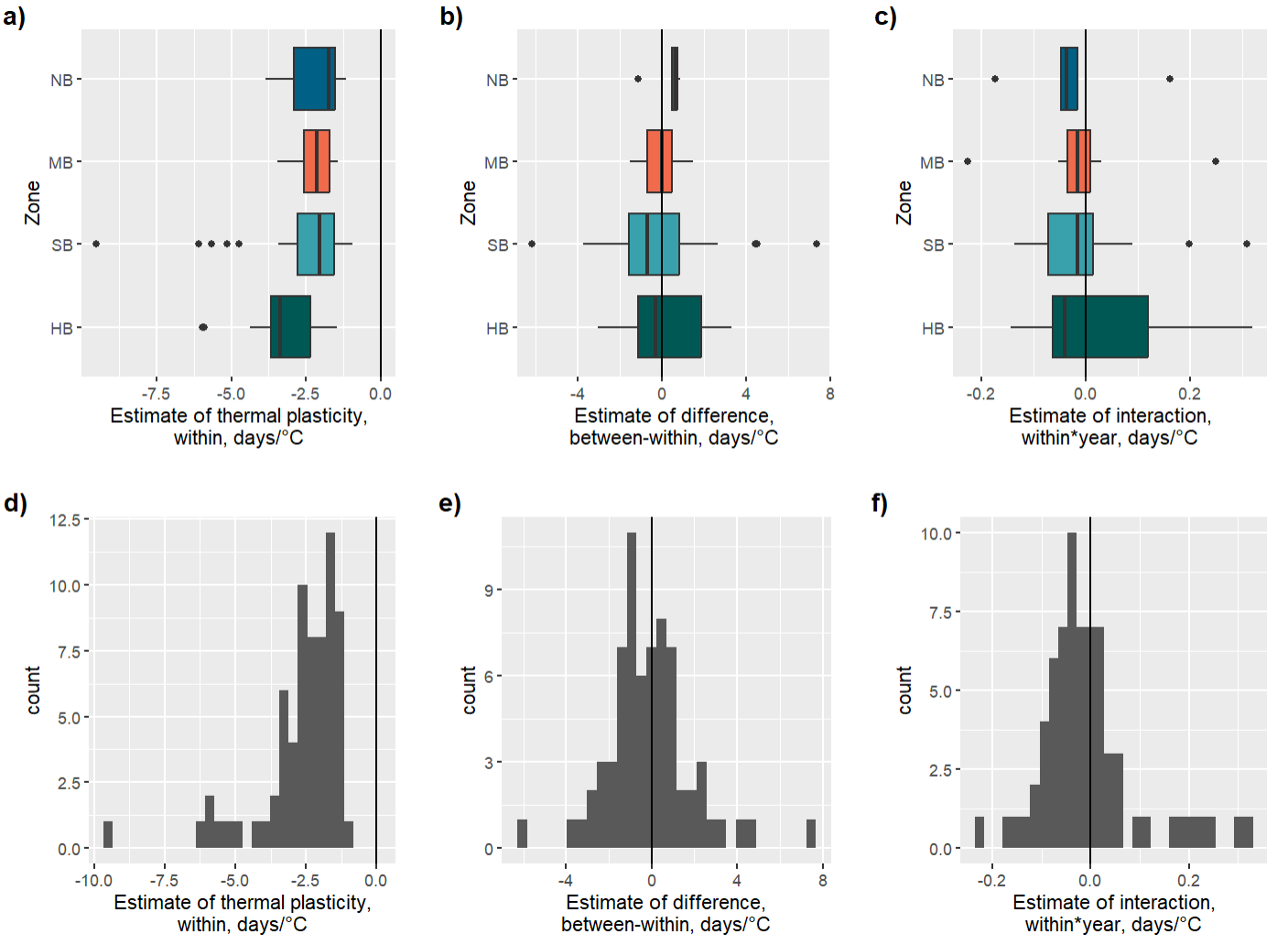

**Figure S6.** Estimates for each species shown as boxplots and histogram for a) and d) average plasticity (within-5-y period effect effect ($\beta_{W})$ parameter from M1), b) and e) difference in reaction norm elevation ($\beta_{B-}\beta_{W}$ from M2), c) and f) change in reaction norm slope (the within-5-y period effect by year interaction term ($\beta_{4}$) from M3. Boxplots organized per bioclimatic zone: HB= hemiboreal zone; SB= southern boreal; MB= middle boreal; NB= northern boreal.

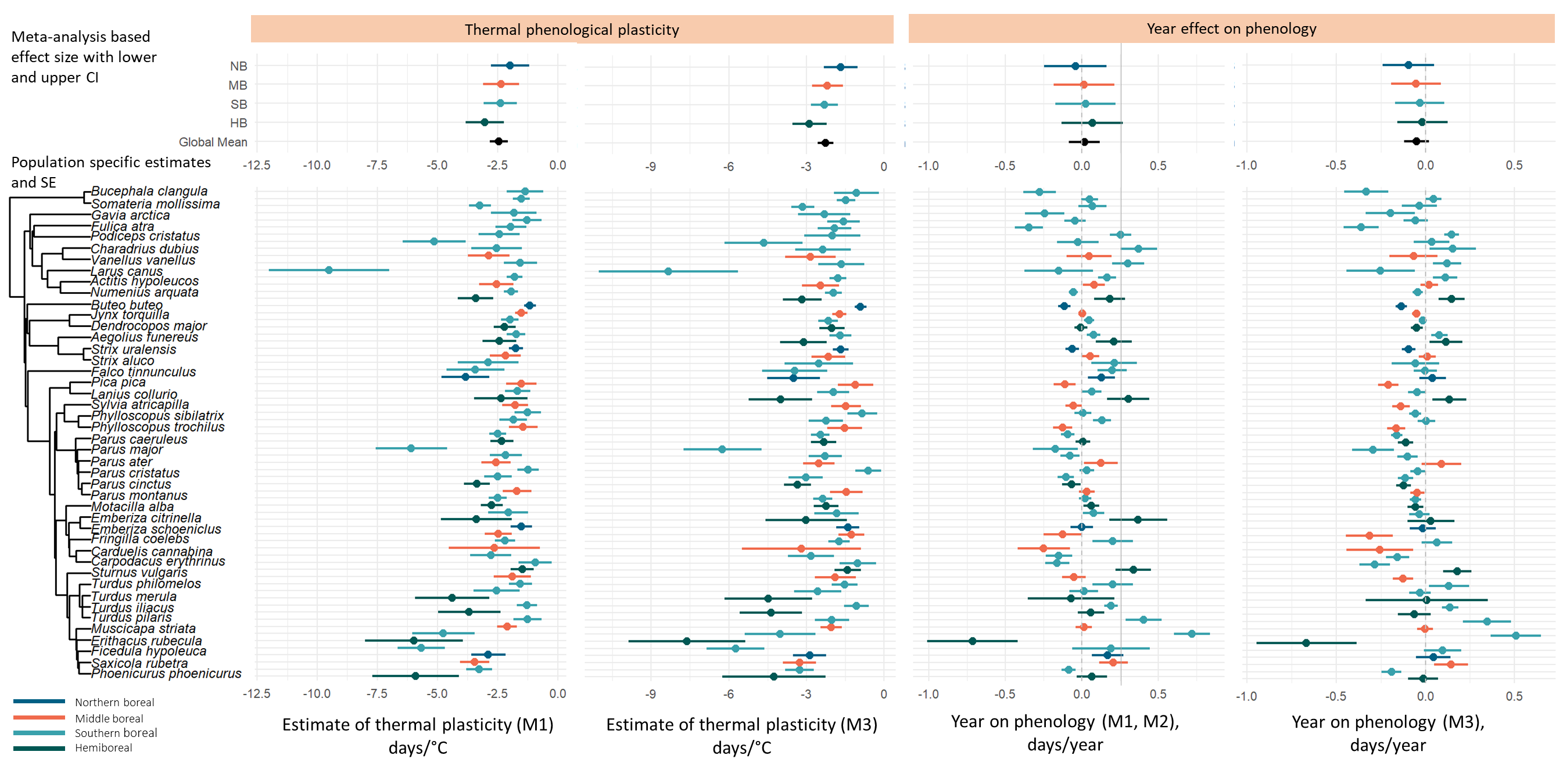
 **Figure S7.** Average plasticity and effect of year. Meta-analysis-based effects sizes with lower and upper confidence intervals per bioclimatic zone and the global mean (upper panels) and population-specific estimates of effect and their standard errors (lower panels) of average plasticity (within-5-y period effect ($\beta_{W})$ parameter from M1 and M3 to the left, and to the right the year effect ($\beta_{3}$ from M1 and M2 (identical) and $\beta_{3}$ from M3), with species ordered by phylogeny and coloured by bioclimatic zone. Exact estimates shown in Tables 1 and S1.

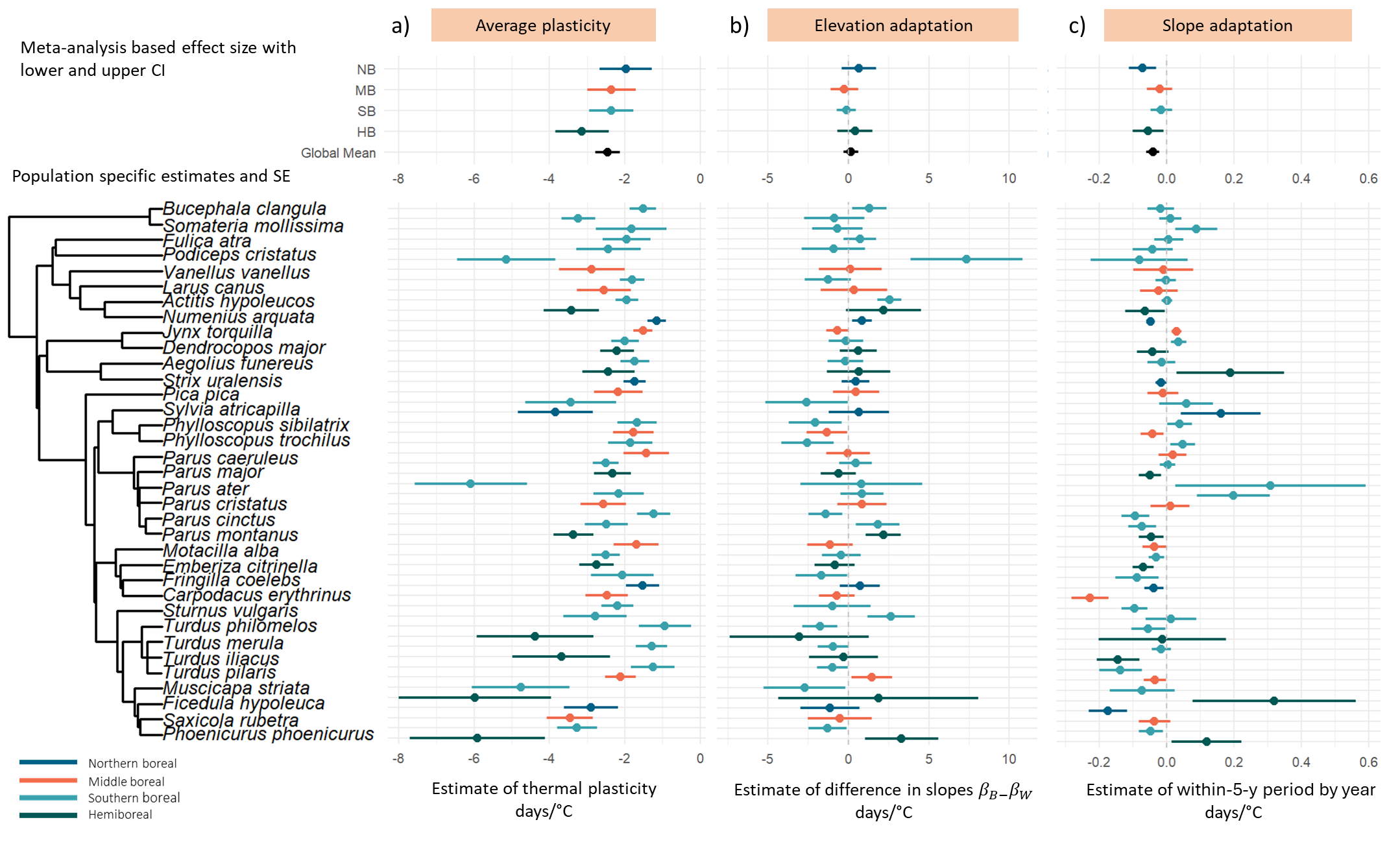
**Figure S8.** Test of average plasticity (leftmost panels) and the hypotheses on *elevation adaptation* (middle panels) and *slope adaptation* (rightmost panels) based on data retained using 0.95 quantile as a threshold for selecting populations with robust cue windows. To identify populations with relatively robust cue windows, we used two approaches (see main text *2.4.* and Texts S2 and S3*)*. For approach 2 we constructed 100 data sets of randomly permutated temperature values. For each of these reshuffled datasets we reran the sliding window approach. For each population, we identified the best model from each year permutation (based on the lowest AIC value). Across the 100 permutated models per population, we then extracted the 0.90 quantile of log likelihood and compared it to the log likelihood of the best model based on correctly paired dates and temperatures. If the log likelihood of the latter was lower than the 0.90 quantile the permutated data we considered the original cue window to be uncertain and discarded the population from the data to be analysed. As a robustness test to the choice of the 0.9 quantile used of the analyses presented in the main text, we also applied an alternative of approach 2 using the 0.95 quantile of log likelihoods, retaining 53 populations of 35 species (16 additional populations removed) with 10 species in the hemiboreal zone, 27 in the southern boreal zone, 11 in the middle boreal zone, and 5 species in the northern boreal zone. The results of the main meta-analyses for the two main hypotheses based on this alternative of using the 0.95 quantile are shown in this figure and in Table S4. Uppermost parts of panels show meta-analysis-based effects sizes with lower and upper confidence intervals per bioclimatic zone and the global mean of a) aim 1 on average plasticity (within-5-y period effect ($\beta_{W})$ parameter from M1); b) aim 2 *on the elevation adaptation hypothesis* (the difference in slopes between versus within-5-y periods ($\beta_{B-}\beta_{W}$) from M2); and c) aim 3 on the *slope adaptation hypothesis* (the within-5-y periods by year interaction term ($\beta_{4}$) from M3). The lower parts of the panels show population-specific estimates and standard errors of respective study questions. Thermal plasticity was somewhat stronger using the 0.95 quantile, but the effect sizes for elevation adaptation and especially slope adaptation were similar.

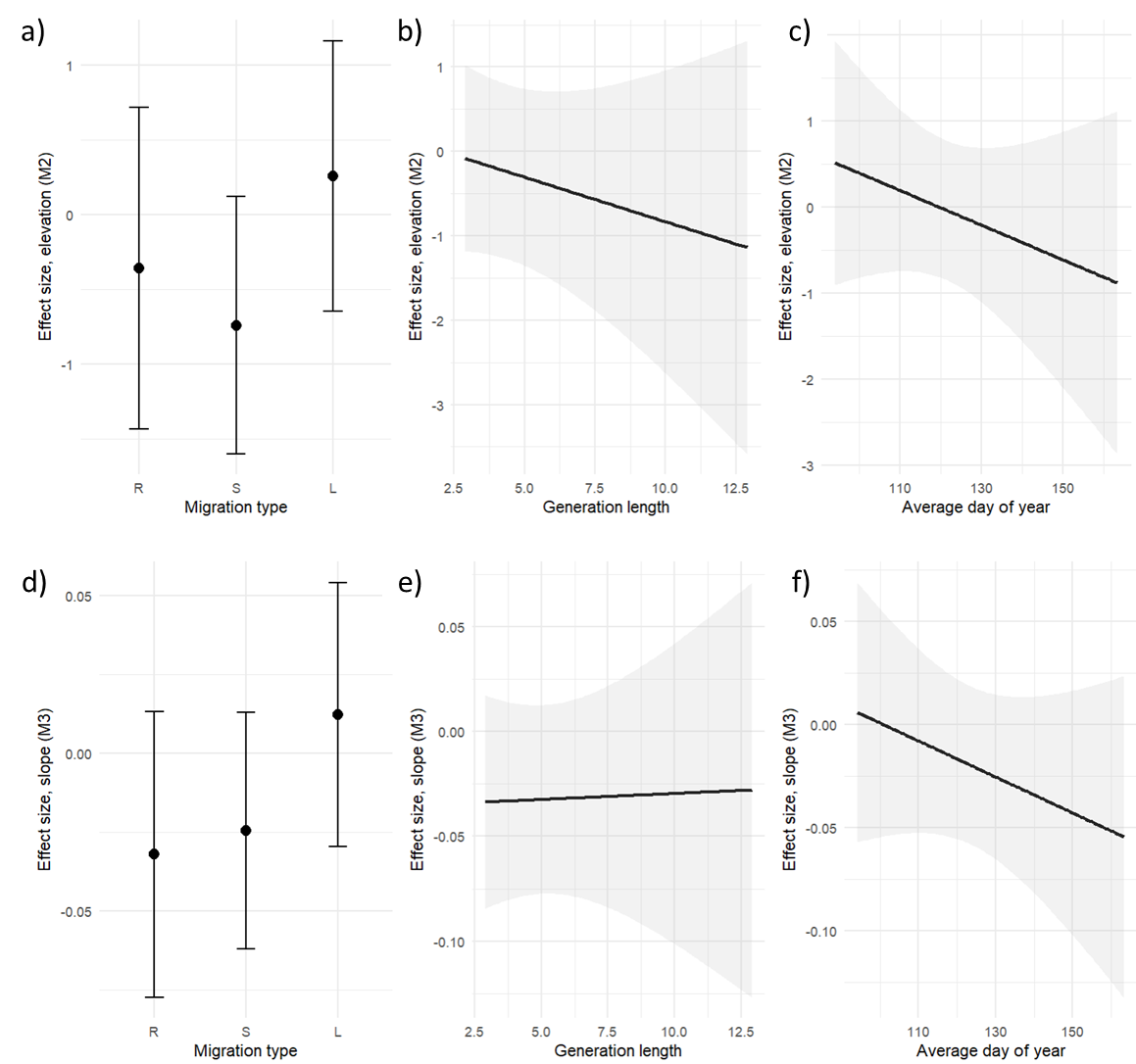

**Figure S9.** Effect size of traits, showing migration type (a and d), generation length (b and e), average FELD (c and f). Panels a)-c) show effects sizes on the meta-analysis-based trait model for aim 2 *on the elevation adaptation hypothesis* (the difference in slopes between versus within-5-y period ($\beta_{B-}\beta_{W}$) from M2), and d)-e) show effects sizes on the meta-analysis-based trait model testing aim 3 on the *slope adaptation hypothesis* (the within-5-y period by year interaction term $\beta_{4}$ from M3). For both aim 2 and 3, the models including traits did not have better explanatory power than a corresponding models without traits as moderators (ΔAIC=2.95 and 3.97 for aim 2 and 3, respectively), and the effect of individual traits was non-significant (Table 3 in main text). The panels show the estimated effect size of each trait when keeping the other traits in the model fixed (generation length and average day of year at their mean value, migration type as Resident, and bioclimatic zone as SB).

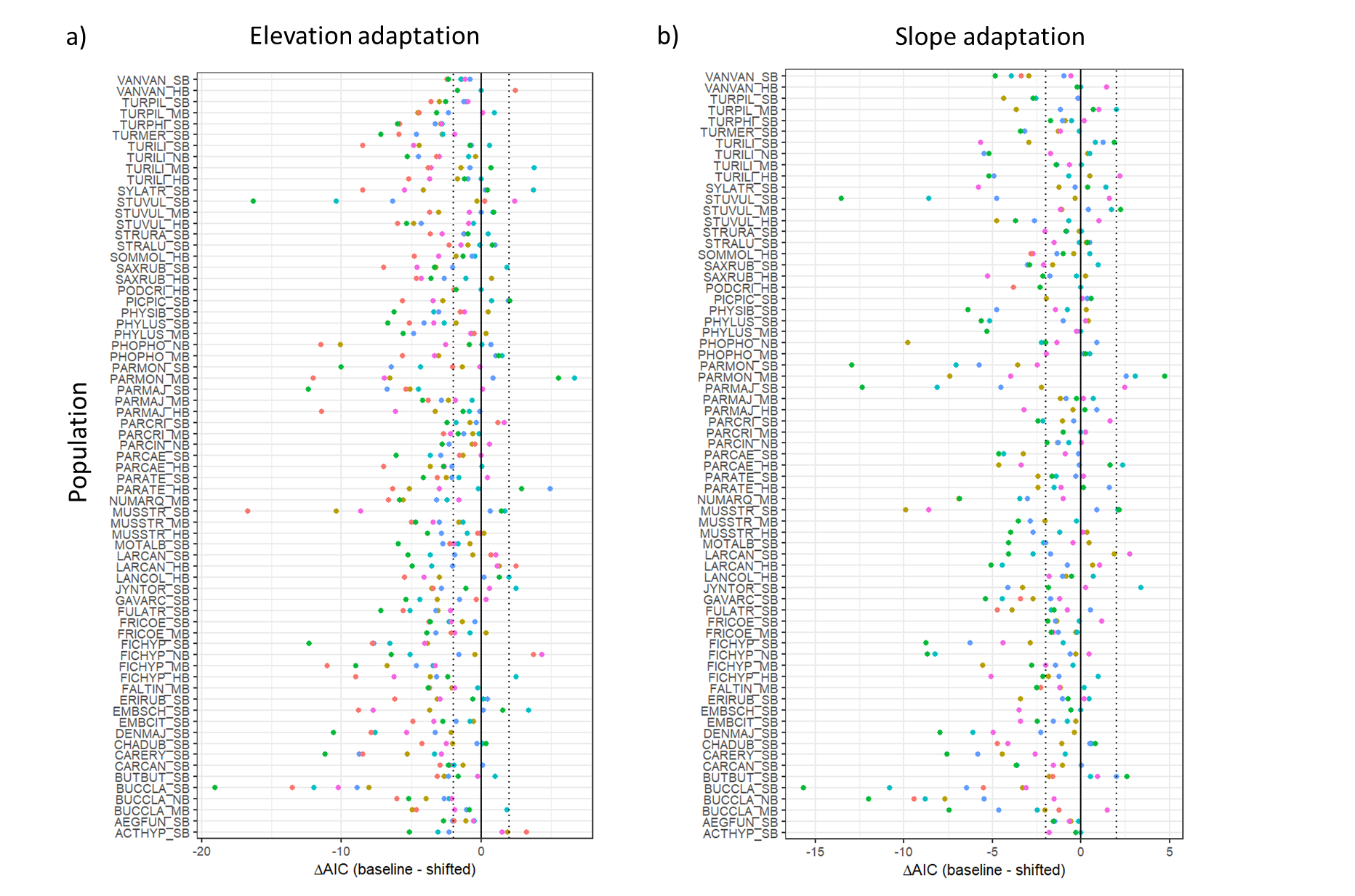

**Figure S10.** Sensitivity test ΔAIC comparison for the potential of shifting cues over time based on a) model testing aim 2 *on the elevation adaptation hypothesis* (the difference in slopes between versus within-5-y period ($\beta_{B-}\beta_{W}$) from M2), and b) model testing aim 3 on the *slope adaptation hypothesis* (the within-5-y period by year interaction term $\beta_{4}$ from M3). Populations are ordered in alphabetical order from bottom to top based on the species name abbreviation (data available in Hällfors et al. (2026)). We divided the data set into two periods (T_1_=1961–1992 and T2=1993–2025) and considered six cases: (1) the climatic cue window used by the species in T_2_ moves to five days earlier, but the window in T_1_ stays the same; (2) the window in T_2_ moves to five days later, T_1_ stays the same; (3) the window in T_2_ stays the same, but the window in T_1_ moves to five days later; (4) the window in T_2_ stays the same, but the window in T_1_ moves to five days earlier; (5) the window in T_2_ moves to five days earlier and the window in T_1_ moves to five days later; (6) the window in T_2_ moves to five days later and the window in T_1_ moves to five days earlier. We found a small proportion where the alternative windows perform better (4.4% and 3.3% for aim 2 and aim 3, respectively, across all alternative models). However, as the frequency of this was low, we assume no general trend in change in cue use and that our approach for the main analyses is robust to such changes. We found no evidence consistent with a shift in the cue window timing, as the AIC values for the six alternative windows tended to be larger and the estimated thermal slopes shallower (Table S1) than in the models based on the best identified cue windows across the complete study time.

**Table S1.** Meta-analysis results of comparing whether average slope estimates are strongest for models based on the overall cue window (defined in section *2.4* in the main text and Texts S2 and S3), or whether any of the cases where the cue window was allowed to change over time provided a more sensitive estimate. The table shows results for models testing aim 2 *on the elevation adaptation hypothesis* (the difference in slopes between versus within-5-y period ($\beta_{B-}\beta_{W}$) from M2), and models testing aim 3 on the *slope adaptation hypothesis* (the within-5-y period by year interaction term $\beta_{4}$ from M3) based on the best identified cue window and six potential cases that we considered. We divided the data set into two periods (T_1_=1961–1992 and T2=1993–2025) and considered six cases: (1) the climatic cue window used by the species in T_2_ moves to five days earlier, but the window in T_1_ stays the same; (2) the window in T_2_ moves to five days later, T_1_ stays the same; (3) the window in T_2_ stays the same, but the window in T_1_ moves to five days later; (4) the window in T_2_ stays the same, but the window in T_1_ moves to five days earlier; (5) the window in T_2_ moves to five days earlier and the window in T_1_ moves to five days later; (6) the window in T_2_ moves to five days later and the window in T_1_ moves to five days earlier. We used a change of five days, as this corresponds with average findings of breeding phenology shifts in birds. σ^2^ indicates the estimated variance of the random effects, in this case species and phylogeny. We found no evidence consistent with a shift in the cue window timing (Fig. S10), as the AIC values for the six alternative windows tended to be larger and the estimated thermal slopes shallower than in the models based on the best identified cue windows across the complete study time. This was confirmed by the meta-analyses presented in this table revealing shallower slopes for mean temperature on phenology.

| **Aim 2 elevation adaptation** | **Estimate** | **SE** | **Zval** | **p-value** | **σ^2^ phylogeny** | **σ^2^ species** |
| --- | --- | --- | --- | --- | --- | --- |
| Best cue window | -3.04 | 0.40 | -7.54 | <0.0001 | 0.49 | 0.00 |
| Case 1 | -2.78 | 0.33 | -8.37 | <0.0001 | 0.25 | 0.00 |
| Case 2 | -2.75 | 0.31 | -8.74 | <0.0001 | 0.22 | 0.00 |
| Case 3 | -2.88 | 0.33 | -8.81 | <0.0001 | 0.22 | 0.00 |
| Case 4 | -2.67 | 0.38 | -7.06 | <0.0001 | 0.41 | 0.00 |
| Case 5 | -2.64 | 0.31 | -8.50 | <0.0001 | 0.17 | 0.00 |
| Case 6 | -2.38 | 0.29 | -8.21 | <0.0001 | 0.17 | 0.02 |
| **Aim 3 slope adaptation** | **Estimate** | **SE** | **Zval** | **p-value** | **σ^2^ phylogeny** | **σ^2^ species** |
| Best cue window | -2.88 | 0.33 | -8.64 | <0.0001 | 0.22 | 0.09 |
| Case 1 | -2.63 | 0.29 | -9.18 | <0.0001 | 0.11 | 0.05 |
| Case 2 | -2.53 | 0.21 | -11.80 | <0.0001 | 0.00 | 0.10 |
| Case 3 | -2.81 | 0.33 | -8.62 | <0.0001 | 0.21 | 0.00 |
| Case 4 | -2.30 | 0.22 | -10.30 | <0.0001 | 0.00 | 0.20 |
| Case 5 | -4.32 | 1.61 | -2.69 | <0.01 | 0.00 | 0.59 |
| Case 6 | -2.17 | 0.21 | -10.40 | <0.0001 | 0.00 | 0.10 |

**Table S2.** Summaries of meta-analysis models for aim 1 on average plasticity (within-5-y period effect ($\beta_{W})$ parameter from M3; within-5-y period effect ($\beta_{W})$ from M1 presented in Table 1 in the main text) and for year effects (year effect ($\beta_{3}$ in M1 and M2 (identical), and $\beta_{3}$ in M3). The results are visualized in Fig S7. The column “Model estimate tested in meta-analysis” indicates which parameter was used a as response in meta-analysis, and from which of the three models (M1, M2, or M3) it was derived. The *BZ* column indicates which zone the estimate refers to, or whether it is the global mean estimate across zones. The estimates for each level of the bioclimatic zone have been adjusted to show the actual effects per level, not the difference to the baseline level. Similarly to the plasticity estimated based on M1, $\beta_{W}$ from M3 indicated and advance of -2.3 days/°C (-2.5, -2) and we found substantial phylogenetic variance in plasticity (σ^2^ = 0.22) and minimal among-species variance (σ^2^ = 0.09). The phylogenetic signal was moderate to strong (0.7). We found no statistically significant effects of year on plasticity ($\beta_{3}$ from M1/2 and M3). σ^2^ indicating the estimated variance of the random effects species and phylogeny, respectively, were 0.0 and 0.4 for $\beta_{3}$ from M1 and M2 and 0.0 and 0.02 for M3, after accounting for fixed effects. The phylogenetic signals were strong for $\beta_{3}$ from both M1/2 and M3 (1.0). An AIC comparison to model versions excluding phylogeny indicated negligible difference for the plasticity model (ΔAIC=-2), the year-model based on effects sizes for $\beta_{3}$ from M1 and M2 (ΔAIC=1.97), and the year-model based on effect sizes from M3 (ΔAIC=1.42). Funnel plots are shown in Fig. S5.

| **Model estimate tested in meta-analysis** | **BZ** | **Estimate** | **lower CI** | **upper CI** | **p-value** |
| --- | --- | --- | --- | --- | --- |
|  | Global mean | **-2.26** | **-2.56** | **-1.96** | **<0.0001** |
| β_W_ from M3 | NB | **-1.67** | **-2.32** | **-1.03** | **<0.0001** |
|  | MB | **-2.19** | **-2.78** | **-1.60** | **<0.0001** |
|  | SB | **-2.30** | **-2.82** | **-1.78** | **<0.0001** |
|  | HB | **-2.88** | **-3.54** | **-2.23** | **<0.0001** |
|  | Global mean | 0.02 | -0.08 | 0.12 | 0.73 |
|  | NB | -0.04 | -0.25 | 0.16 | 0.70 |
| β_3_ from M1 and M2 | MB | 0.02 | -0.18 | 0.22 | 0.87 |
|  | SB | 0.03 | -0.17 | 0.22 | 0.80 |
|  | HB | 0.07 | -0.13 | 0.27 | 0.49 |
|  | Global mean | -0.05 | -0.12 | 0.02 | 0.17 |
|  | NB | -0.10 | -0.24 | 0.05 | 0.20 |
| β_3_ from M3 | MB | -0.05 | -0.19 | 0.09 | 0.46 |
|  | SB | -0.03 | -0.17 | 0.11 | 0.66 |
|  | HB | -0.02 | -0.16 | 0.12 | 0.82 |

**Table S3. Population specific results for study aim 1-3.** Aim 1 on average plasticity (within-5-y period effect ($\beta_{W})$ parameter from M1; aim 2 *on the elevation adaptation hypothesis* (the difference in slopes between versus within-5-y period ($\beta_{B-}\beta_{W}$) from M2); and aim 3 on the *slope adaptation hypothesis* (the within-5-y period by year interaction term ($\beta_{4}$) from M3). NB= Northern boreal; MB= Middle boreal; SB= Southern boreal; HB= hemiboreal; SE= standard error. Upper and lower CI indicate the upper and lower bounds of 95% confidence intervals and are shown in bold when excluding zero.

|  |  | **Aim 1: Average plasticity** | | | | **Aim 2: Elevation adaptation** | | | | **Aim 3: Slope adaptation** | | | |
| --- | --- | --- | --- | --- | --- | --- | --- | --- | --- | --- | --- | --- | --- |
| **Species** | **Bioclimatic Zone** | **Estimate** | **Standard error** | **lower CI** | **upper CI** | **Estimate** | **Standard error** | **lower CI** | **upper CI** | **Estimate** | **Standard error** | **lower CI** | **upper CI** |
| Actitis hypoleucos | SB | -1.26 | 0.58 | **-2.39** | **-0.13** | -0.98 | 0.96 | -2.87 | 0.91 | -0.14 | 0.06 | **-0.26** | **-0.01** |
| Aegolius funereus | SB | -1.95 | 0.64 | **-3.20** | **-0.70** | 0.72 | 1.03 | -1.30 | 2.73 | 0.01 | 0.04 | -0.08 | 0.09 |
| Bucephala clangula | MB | -3.46 | 0.61 | **-4.66** | **-2.25** | -0.53 | 1.98 | -4.40 | 3.35 | -0.04 | 0.05 | -0.13 | 0.06 |
| Bucephala clangula | NB | -2.89 | 0.71 | **-4.29** | **-1.50** | -1.13 | 1.84 | -4.74 | 2.48 | -0.17 | 0.06 | **-0.28** | **-0.06** |
| Bucephala clangula | SB | -3.26 | 0.53 | **-4.31** | **-2.22** | -1.29 | 1.20 | -3.63 | 1.06 | -0.05 | 0.04 | -0.12 | 0.02 |
| Buteo buteo | SB | -1.36 | 0.76 | -2.85 | 0.12 | -1.59 | 1.35 | -4.23 | 1.05 | -0.06 | 0.10 | -0.25 | 0.13 |
| Carduelis cannabina | SB | -9.51 | 2.50 | **-14.41** | **-4.61** | 4.50 | 6.11 | -7.48 | 16.48 | -0.12 | 0.13 | -0.38 | 0.14 |
| Carpodacus erythrinus | SB | -1.80 | 0.33 | **-2.45** | **-1.16** | -1.27 | 1.45 | -4.11 | 1.57 | 0.00 | 0.03 | -0.06 | 0.06 |
| Charadrius dubius | SB | -1.55 | 0.48 | **-2.50** | **-0.61** | 0.11 | 1.64 | -3.10 | 3.33 | 0.01 | 0.06 | -0.11 | 0.14 |
| Dendrocopos major | SB | -3.24 | 0.45 | **-4.12** | **-2.36** | -0.87 | 1.87 | -4.53 | 2.79 | 0.01 | 0.03 | -0.05 | 0.08 |
| Emberiza citrinella | SB | -5.14 | 1.31 | **-7.70** | **-2.58** | 7.34 | 3.48 | **0.51** | **14.16** | -0.08 | 0.14 | -0.36 | 0.20 |
| Emberiza schoeniclus | SB | -2.54 | 1.04 | **-4.59** | **-0.50** | -2.23 | 1.90 | -5.95 | 1.50 | -0.06 | 0.13 | -0.31 | 0.19 |
| Erithacus rubecula | SB | -2.90 | 1.26 | **-5.36** | **-0.44** | -6.19 | 3.94 | -13.90 | 1.53 | -0.08 | 0.08 | -0.23 | 0.07 |
| Falco tinnunculus | MB | -1.89 | 0.77 | **-3.40** | **-0.38** | 0.01 | 1.55 | -3.02 | 3.04 | 0.00 | 0.05 | -0.09 | 0.10 |
| Ficedula hypoleuca | HB | -2.21 | 0.45 | **-3.08** | **-1.33** | 0.61 | 1.15 | -1.64 | 2.85 | -0.04 | 0.05 | -0.13 | 0.05 |
| Ficedula hypoleuca | MB | -1.52 | 0.26 | **-2.02** | **-1.01** | -0.68 | 0.70 | -2.05 | 0.69 | 0.03 | 0.01 | **0.00** | **0.06** |
| Ficedula hypoleuca | NB | -1.16 | 0.24 | **-1.63** | **-0.68** | 0.85 | 0.63 | -0.37 | 2.08 | -0.05 | 0.01 | **-0.07** | **-0.02** |
| Ficedula hypoleuca | SB | -1.99 | 0.36 | **-2.70** | **-1.28** | -0.15 | 1.07 | -2.24 | 1.94 | 0.04 | 0.02 | -0.01 | 0.08 |
| Fringilla coelebs | MB | -2.87 | 0.87 | **-4.58** | **-1.16** | 0.12 | 1.96 | -3.72 | 3.95 | -0.01 | 0.09 | -0.18 | 0.16 |
| Fringilla coelebs | SB | -1.55 | 0.70 | **-2.92** | **-0.19** | -3.74 | 2.48 | -8.61 | 1.13 | 0.01 | 0.05 | -0.08 | 0.10 |
| Fulica atra | SB | -4.76 | 1.30 | **-7.30** | **-2.21** | -2.71 | 2.54 | -7.68 | 2.26 | -0.07 | 0.10 | -0.26 | 0.12 |
| Gavia arctica | SB | -5.67 | 0.99 | **-7.60** | **-3.74** | 4.40 | 3.02 | -1.51 | 10.31 | 0.02 | 0.13 | -0.23 | 0.28 |
| Jynx torquilla | SB | -1.52 | 0.35 | **-2.20** | **-0.83** | 1.31 | 1.09 | -0.82 | 3.43 | -0.02 | 0.04 | -0.10 | 0.06 |
| Lanius collurio | HB | -1.48 | 0.48 | **-2.42** | **-0.53** | -1.16 | 1.54 | -4.17 | 1.86 | 0.00 | 0.04 | -0.09 | 0.08 |
| Larus canus | HB | -3.68 | 1.30 | **-6.23** | **-1.14** | -0.31 | 2.14 | -4.51 | 3.89 | -0.14 | 0.06 | **-0.27** | **-0.02** |
| Larus canus | SB | -1.29 | 0.42 | **-2.11** | **-0.47** | -0.94 | 0.96 | -2.82 | 0.94 | -0.02 | 0.03 | -0.07 | 0.04 |
| Motacilla alba | SB | -2.43 | 0.85 | **-4.11** | **-0.76** | -0.92 | 1.97 | -4.79 | 2.96 | -0.04 | 0.06 | -0.16 | 0.08 |
| Muscicapa striata | HB | -3.42 | 0.73 | **-4.85** | **-1.99** | 2.19 | 2.34 | -2.41 | 6.78 | -0.06 | 0.06 | -0.18 | 0.05 |
| Muscicapa striata | MB | -2.55 | 0.71 | **-3.95** | **-1.15** | 0.36 | 2.07 | -3.69 | 4.42 | -0.02 | 0.06 | -0.13 | 0.09 |
| Muscicapa striata | SB | -1.95 | 0.30 | **-2.53** | **-1.36** | 2.56 | 0.74 | **1.10** | **4.02** | 0.00 | 0.02 | -0.03 | 0.03 |
| Numenius arquata | MB | -2.11 | 0.41 | **-2.91** | **-1.31** | 1.46 | 1.28 | -1.04 | 3.97 | -0.03 | 0.03 | -0.10 | 0.03 |
| Parus ater | HB | -3.39 | 1.47 | **-6.27** | **-0.51** | -2.48 | 2.77 | -7.90 | 2.94 | -0.09 | 0.17 | -0.42 | 0.23 |
| Parus ater | SB | -2.06 | 0.83 | **-3.68** | **-0.44** | -1.66 | 1.60 | -4.80 | 1.49 | -0.09 | 0.06 | -0.21 | 0.04 |
| Parus caeruleus | HB | -3.36 | 0.53 | **-4.41** | **-2.32** | 2.17 | 1.11 | 0.00 | 4.34 | -0.05 | 0.04 | -0.12 | 0.03 |
| Parus caeruleus | SB | -2.49 | 0.58 | **-3.62** | **-1.36** | 1.83 | 1.35 | -0.81 | 4.46 | -0.07 | 0.04 | -0.15 | 0.01 |
| Parus cinctus | NB | -1.52 | 0.44 | **-2.38** | **-0.66** | 0.72 | 1.25 | -1.73 | 3.17 | -0.04 | 0.03 | -0.09 | 0.02 |
| Parus cristatus | MB | -2.63 | 1.90 | -6.36 | 1.09 | 0.54 | 2.08 | -3.53 | 4.62 | 0.25 | 0.36 | -0.45 | 0.95 |
| Parus cristatus | SB | -2.79 | 0.84 | **-4.44** | **-1.13** | 2.65 | 1.47 | -0.23 | 5.53 | 0.01 | 0.08 | -0.13 | 0.16 |
| Parus major | HB | -2.75 | 0.45 | **-3.64** | **-1.86** | -0.85 | 1.24 | -3.27 | 1.57 | -0.07 | 0.03 | **-0.13** | **-0.01** |
| Parus major | MB | -1.70 | 0.60 | **-2.86** | **-0.53** | -1.14 | 1.41 | -3.89 | 1.62 | -0.04 | 0.03 | -0.10 | 0.03 |
| Parus major | SB | -2.50 | 0.37 | **-3.24** | **-1.77** | -0.44 | 1.20 | -2.80 | 1.92 | -0.03 | 0.02 | -0.07 | 0.01 |
| Parus montanus | MB | -2.48 | 0.56 | **-3.57** | **-1.38** | -0.71 | 1.11 | -2.88 | 1.46 | -0.23 | 0.06 | **-0.34** | **-0.12** |
| Parus montanus | SB | -2.19 | 0.42 | **-3.02** | **-1.37** | -0.99 | 2.39 | -5.67 | 3.69 | -0.09 | 0.04 | **-0.17** | **-0.02** |
| Phoenicurus phoenicurus | MB | -2.17 | 0.64 | **-3.43** | **-0.92** | 0.48 | 1.43 | -2.33 | 3.29 | -0.01 | 0.05 | -0.10 | 0.08 |
| Phoenicurus phoenicurus | NB | -1.74 | 0.29 | **-2.32** | **-1.16** | 0.46 | 0.86 | -1.22 | 2.15 | -0.02 | 0.02 | -0.05 | 0.02 |
| Phylloscopus sibilatrix | SB | -2.16 | 0.67 | **-3.47** | **-0.85** | 0.85 | 1.35 | -1.78 | 3.49 | 0.20 | 0.11 | -0.01 | 0.41 |
| Phylloscopus trochilus | MB | -2.57 | 0.60 | **-3.75** | **-1.39** | 0.86 | 1.53 | -2.14 | 3.87 | 0.01 | 0.06 | -0.10 | 0.13 |
| Phylloscopus trochilus | SB | -1.23 | 0.44 | **-2.09** | **-0.38** | -1.42 | 1.06 | -3.49 | 0.65 | -0.09 | 0.04 | **-0.17** | **-0.01** |
| Pica pica | SB | -0.94 | 0.69 | -2.30 | 0.42 | -1.76 | 1.10 | -3.91 | 0.39 | -0.05 | 0.05 | -0.15 | 0.04 |
| Podiceps cristatus | HB | -5.98 | 2.02 | **-9.95** | **-2.01** | 1.87 | 6.22 | -10.32 | 14.05 | 0.32 | 0.24 | -0.16 | 0.80 |
| Saxicola rubetra | HB | -2.43 | 0.70 | **-3.80** | **-1.07** | 0.65 | 1.96 | -3.20 | 4.49 | 0.19 | 0.16 | -0.12 | 0.50 |
| Saxicola rubetra | SB | -1.73 | 0.39 | **-2.49** | **-0.97** | -0.18 | 1.11 | -2.35 | 1.99 | -0.01 | 0.04 | -0.10 | 0.07 |
| Somateria mollissima | HB | -5.91 | 1.79 | **-9.43** | **-2.40** | 3.30 | 2.28 | -1.17 | 7.78 | 0.12 | 0.10 | -0.08 | 0.32 |
| Strix aluco | SB | -1.28 | 0.61 | **-2.49** | **-0.08** | -0.11 | 1.04 | -2.14 | 1.92 | 0.06 | 0.03 | 0.00 | 0.12 |
| Strix uralensis | SB | -1.83 | 0.94 | -3.67 | 0.02 | -0.69 | 1.57 | -3.76 | 2.38 | 0.09 | 0.06 | -0.03 | 0.21 |
| Sturnus vulgaris | HB | -2.32 | 0.49 | **-3.28** | **-1.37** | -0.62 | 1.10 | -2.78 | 1.54 | -0.05 | 0.03 | -0.11 | 0.02 |
| Sturnus vulgaris | MB | -1.44 | 0.60 | **-2.62** | **-0.25** | -0.01 | 1.35 | -2.66 | 2.63 | 0.02 | 0.04 | -0.06 | 0.10 |
| Sturnus vulgaris | SB | -2.51 | 0.35 | **-3.19** | **-1.82** | 0.45 | 1.01 | -1.53 | 2.43 | 0.00 | 0.02 | -0.04 | 0.05 |
| Sylvia atricapilla | SB | -6.09 | 1.49 | **-9.01** | **-3.16** | 0.80 | 3.79 | -6.63 | 8.23 | 0.31 | 0.28 | -0.24 | 0.86 |
| Turdus iliacus | HB | -2.37 | 1.10 | **-4.53** | **-0.21** | -1.41 | 2.35 | -6.02 | 3.20 | 0.23 | 0.09 | **0.06** | **0.40** |
| Turdus iliacus | MB | -1.52 | 0.63 | **-2.75** | **-0.28** | -1.54 | 1.72 | -4.92 | 1.84 | -0.05 | 0.04 | -0.13 | 0.02 |
| Turdus iliacus | NB | -3.84 | 1.00 | **-5.80** | **-1.88** | 0.65 | 1.88 | -3.03 | 4.33 | 0.16 | 0.12 | -0.07 | 0.39 |
| Turdus iliacus | SB | -1.68 | 0.52 | **-2.70** | **-0.65** | -2.06 | 1.63 | -5.27 | 1.14 | 0.04 | 0.04 | -0.03 | 0.11 |
| Turdus merula | SB | -3.43 | 1.20 | **-5.79** | **-1.07** | -2.59 | 2.56 | -7.60 | 2.43 | 0.06 | 0.08 | -0.10 | 0.21 |
| Turdus philomelos | SB | -1.85 | 0.58 | **-2.99** | **-0.72** | -2.54 | 1.63 | -5.73 | 0.64 | 0.05 | 0.04 | -0.02 | 0.12 |
| Turdus pilaris | MB | -1.77 | 0.54 | **-2.82** | **-0.71** | -1.32 | 1.26 | -3.78 | 1.15 | -0.04 | 0.03 | -0.11 | 0.03 |
| Turdus pilaris | SB | -1.25 | 0.55 | **-2.33** | **-0.16** | -0.71 | 1.45 | -3.54 | 2.13 | -0.07 | 0.03 | **-0.14** | **-0.01** |
| Vanellus vanellus | HB | -4.39 | 1.55 | **-7.42** | **-1.35** | -3.06 | 4.32 | -11.53 | 5.41 | -0.01 | 0.19 | -0.38 | 0.36 |
| Vanellus vanellus | SB | -2.55 | 0.95 | **-4.41** | **-0.69** | 1.16 | 2.45 | -3.64 | 5.95 | -0.11 | 0.05 | **-0.22** | **0.00** |

**Table S4. Summaries of meta-analysis models for** average plasticity, and the hypotheses on *elevation adaptation* and *slope adaptation* based on data retained using 0.95 quantile as a threshold for selecting populations with robust cue windows. To identify populations with relatively robust cue windows, we used two approaches (see main text *2.4.*and Texts S2 and S3). The results of the main meta-analyses for the two main hypotheses based on this alternative of using the 0.95 quantile are visualised in Fig. S8. This table presents the meta-analysis-based effects sizes with lower and upper confidence intervals per bioclimatic zone and the global mean of aim 1 on average plasticity (within-5-y period effect $\beta_{W}$ parameter from M1), aim 2 *on the elevation adaptation hypothesis* (the difference in slopes between versus within-5-y period $\beta_{B-}\beta_{W}$ from M2), and aim 3 on the *slope adaptation hypothesis* (the within-5-y period by year interaction term $\beta_{4}$ from M3). Thermal plasticity was somewhat stronger using the 0.95 quantile, but the effect sizes for elevation adaptation and especially slope adaptation were similar. The column “Study question and model estimate tested in meta-analysis” indicates which study question and model estimate was used a as response in meta-analysis, and from which of the three models it was derived. The *BZ* column indicates which zone the estimate refers to, or whether it is the global mean estimate across zones. The estimates for each level of the bioclimatic zone have been adjusted to show the actual effects per level, not the difference to the baseline level. Statistically significant effects in bold. For aim 1, σ^2^ indicating the estimated variance of the random effects species and phylogeny, respectively, were 0 and 0.29, after accounting for fixed effects. The phylogenetic signal was strong (1.0). The corresponding σ^2^ for aim 2 and 3, respectively, were 0.55 and, and 0 and 0. The phylogenetic signals were low for aim 2 (0.0) and moderate for aim 3 (0.57). An AIC comparison to a model version excluding phylogeny indicated a negligible difference between the models with ΔAIC=-1.2, -2, and -2 for aims 1, 2, and 3, respectively, with a negative AIC value indicating a better fit for the model without a phylogeny. Compared to the main results based on the 0.9 quantile, the effect sizes were identical in direction and similar in magnitude. Thermal plasticity somewhat stronger using the 0.95 quantile, but the effect sizes for elevation adaptation and especially slope adaptation were similar.

| **Aim and model estimate tested in meta-analysis** | **BZ** | **Estimate** | **lower CI** | **upper CI** | **p-value** |
| --- | --- | --- | --- | --- | --- |
| Aim 1: plasticity | Global mean | **-2.46** | **-2.79** | **-2.13** | **<0.0001** |
|  | NB | **-1.98** | **-2.67** | **-1.29** | **<0.0001** |
| β_w_ from M1 | MB | **-2.35** | **-3.00** | **-1.71** | **<0.0001** |
|  | SB | **-2.37** | **-2.95** | **-1.78** | **<0.0001** |
|  | HB | **-3.13** | **-3.85** | **-2.42** | **<0.0001** |
| Aim 2: elevation adaptation | Global mean | 0.18 | -0.29 | 0.64 | 0.45 |
|  | NB | 0.66 | -0.40 | 1.72 | 0.22 |
| β_B_ – β_W_ from M2 | MB | -0.24 | -1.12 | 0.63 | 0.58 |
|  | SB | -0.12 | -0.70 | 0.45 | 0.68 |
|  | HB | 0.41 | -0.68 | 1.50 | 0.46 |
| Aim 3: slope adaptation | Global mean | **-0.04** | **-0.06** | **-0.02** | **<0.0001** |
|  | NB | **-0.07** | **-0.11** | **-0.03** | **<0.001** |
| β_4_ from M3 | MB | -0.02 | -0.06 | 0.02 | 0.16 |
|  | SB | -0.02 | -0.05 | 0.02 | 0.20 |
|  | HB | **-0.06** | **-0.10** | **-0.009** | **<0.05** |

**Text S1. Converting accuracy levels into measurement error variance**

The accuracy level defined by Kluen et al. (2017; Table on page 7 in the supplementary materials, an integer of 1…6 where larger values indicate a closer temporal accuracy) was converted into a measurement error variance around the estimated day of hatching. This was done by taking the variance of a normal distribution where the 99.5th percentile corresponds to the uncertainty in FELD (expressed as the integer accuracy levels from Kluen et al. 2017). This yielded a link between the accuracy level and variance, with a larger date imprecision (lower accuracy) producing a larger variance (accuracy 1 = a variance of 15.07; acc. 2 = var. 3.77; acc. 3 = var. 1.36; acc. 4 = var. 0.60; acc. 5 = var 0.15) and thus, by applying 1/variance, a lower weight to be used in the models (see 2.6 Testing for shifts in response norms). In the data, there were no observations with an accuracy level of 6 (visit on exact time of egg-laying) but for consistency, we defined the variance for such a case to correspond to a conservative ±0.5 day interval around FELD and a corresponding variance of 0.04.

**Text S2. Sliding window approach for cue window identification**

Before estimating temperature-phenology reaction norms, it is important to identify (i) the environmental cues that a population’s FELD is most responsive to and (ii) whether FELD is temperature sensitive. Here, we use a sliding window approach (e.g. Brommer et al., 2008; van de Pol et al., 2016), applying the approach separately for each population (i.e., species in a bioclimatic zone). We fitted sliding window models relating the FELD of each population across the whole study period as a function of the average daily temperature in that zone during different seasonal time periods, ranging from a start on day of the year 60 (March 1^st^ in non-leap years) and an end on the day of the year that corresponded to the mean FELD of the population. We restricted the starting day to day 60, as we anticipate that birds will respond primarily to spring temperatures (the average start of the growing season in Finland in the end of April; Aalto et al., 2022), and this reduces the extent of multiple-testing. For the study bird species in Finland, spring nesting typically starts from March onwards (Lehikoinen et al., 2022; Lehikoinen & Vähätalo, 2000). We allowed the windows to vary in duration between 30–100 days in 1-day increments.

Models included *year* as a fixed effect to detrend the analysis (Keogan et al., 2018; Macphie et al., 2025), and *year* and *site* as random terms to account for temporal and spatial structure in the observations, respectively. We created the site variable by rounding the northing (latitude) coordinates to two significant figures and the easting (longitude) to one significant digit, resulting in grid cells of 10 km latitude x 100 km longitude, 55 of which had nesting observation data. We opted for rectangular grids as we anticipate that temperature and phenology will vary more across latitudes than across longitudes. We introduced weights to FELD observations by using the *accuracy* of the FELD estimate (measurement error variance around the estimated FELD; Text S1).

To identify the cue window during which the average temperature best explained variation in average FELD for a population we used AIC (Akaike’s Information Criterion; Burnham & Anderson, 2004), obtained from mixed models that used maximum likelihood (*lmer* in the lme4 package in R; Bates et al., 2015).

**Text S3.** **Removing populations with uncertain cue windows**

Even under a null scenario where a species is not responsive to a thermal cue during the spring, the sliding window approach will still find a window that yields the lowest AIC (Bailey & Pol, 2016). To minimise the potential for downstream analyses to include populations for which there is little evidence for a relationship between temperature and phenology, we adopted two approaches to identify such populations and exclude them. (i) Populations with uncertain window position – where windows within 2 AIC of the best window differed in starting or ending day by more than 30 days compared to the best window (2 populations and 1 species removed, 139 populations and 76 species retained). (ii) Non-rejection of permutation based null – we followed Macphie et al. (2025) in constructing 100 data sets for each species in which the temperature values were permuted across years (rather than across all observations). For each of these permuted datasets we reran the sliding window approach and identified the best model from each permutation (based on the lowest AIC value). Then we assessed whether the log likelihood of the data given the true temperature data exceeded the 90^th^ quantile of 100 log likelihoods obtained across the permutations. Where it did not, we considered the original cue window to lack support (non-rejection of the permutation null,70 additional populations and 32 species were removed).

Adding the two filters above resulted in retention of 69 populations represented by 44 species (49% of our original number of populations in the data set). For the analyses, we thus ended up with 13 species in the hemiboreal zone, 37 in the southern boreal zone, 14 in the middle boreal zone, and five in the northern boreal zone (Fig. 2 in main text; Fig. S1 e-f). As a robustness test, we also applied an alternative approach (ii), using the 95^th^ quantile of log likelihoods, which retained 53 populations of 35 species (37% of populations).

**Text S4.** **Robustness test of potential shifts in cue timing**

As our correlation-based inference approach is sensitive to the assumption that we have identified the causal cue for phenology (Tansey et al., 2017), we also examine whether there are shifts in FELD over time that are not captured by temperature effects (testing the effect of year, described in Section *2.6*. in the main text), and whether the timing of the temperature cues’ themselves have shifted over time.

It is possible that the cue that the populations use for initiating their breeding phenology may have shifted earlier or later over time, or that the correlation between temperature and another proximate environmental cue has changed over time, leading to a shift in the seasonal period of temperature sensitivity (Bonamour et al., 2019; Simmonds et al., 2019). To examine whether our key findings were robust to this possibility of shifting cue timing, we performed sensitivity analyses testing whether the thermal cue windows differed for the analysed populations between the first and second half of the study period. We divided the data set into two periods (T_1_=1961–1992 and T2=1993–2025) and considered six cases: (i) the climatic cue window used by the population in T_2_ moves to five days earlier, but the window in T_1_ stays the same (i.e. equaling the best window estimated in Section 2.4); (ii) the window in T_2_ moves to five days later, T_1_ stays the same; (iii) the window in T_2_ stays the same, but the window in T_1_ moves to five days later; (iv) the window in T_2_ stays the same, but the window in T_1_ moves to five days earlier; (v) the window in T_2_ moves to five days earlier and the window in T_1_ moves to five days later; (vi) the window in T_2_ moves to five days later and the window in T_1_ moves to five days earlier. We hold the magnitude of shift constant to minimize multiple-testing (Bailey & van de Pol, 2016). We compared the AIC values and estimated slopes of the six alternative models with those of the best cue window. We calculated the AIC based on the model loglikelihood with the degrees of freedom equal to the model degrees of freedom plus four for the alternative models and plus two for the original ones, to account for the additional start and end dates (following Phillimore et al., 2016).

In addition to visual inspection of the differences and the AIC comparison between the six cases, we used a meta-analysis to test whether average slope estimates were strongest for models based on the overall cue window (defined in Section *2.4* and Text S2), or whether any of the cases where the cue window is allowed to change over time provides a more sensitive estimate. Here, we compared the temperature slope estimates of the shifting cue windows to the best cue window identified in Section *2.4* in the main text and Text S2, i.e. we included both estimates as responses and included a moderator for cue type (single cue, versus shifting cue) and a random effect for each species by bioclimate zone.

We found no evidence consistent with a shift in the cue window timing (Fig. S10, Table S1), as the AIC values for the six alternative windows tended to be larger for the models based on the best identified cue windows across the complete study time, with only 4.4% and 3.3% for aim 2 and aim 3, respectively, having ΔAIC>2 across all alternative models (Fig. S10). This was confirmed by meta-analyses on the estimates from models based on shifting cue, which revealed shallower slopes for mean temperature on phenology compared to models based on the best identified cue windows across the complete study time (Table S1).

The average effect of year ($\beta_{3})$ on the change in phenology indicated no significant independent effect of year on FELD (Table S2), neither when included as an independent effect (M1 and M2: 0.02 days/year, 95% CI: -0.08, 0.12;) nor when it was allowed to interact with short-term phenological slopes (M3: -0.05 days/year, 95% CI: -0.12, 0.02) (Table S2). We found a small phylogenetic effect (σ2 = 0.04 and 0.02 for M1and M2, and M3, respectively) and no among-species variance in the meta-analysis models testing year-effects. This was supported by an AIC comparison to model version excluding phylogeny, indicating a small difference between the models in favour of the model including phylogeny (ΔAIC = 4.54 and -4.05 for M1 and M2, and M3, respectively, with a negative AIC value indicating a better fit for the model without a phylogeny).

**Text S5. Methodological Considerations and Limitations**

We performed a series of robustness checks and sensitivity analyses to evaluate the reliability of cue window identification, the stability of thermal reaction norm estimates, and potential sources of statistical bias. These included filtering populations with uncertain cue windows, permutation tests, alternative quantile thresholds for cue identification, and two-period sensitivity analyses to test for temporal shifts in cue timing. Collectively, these analyses indicate that the identified thermal cue windows are generally robust, unlikely to have shifted substantially over time, and not driven by changes in temperature variability or sampling structure. At the same time, our approach entails important methodological considerations, including statistical coupling between within- and between-population effects (see below; Phillimore et al., 2010; Westneat et al., 2020), uncertainty in slope estimates arising from sliding window selection, and biases introduced by retaining populations with strong temperature signals. While these caveats do not appear to undermine the main conclusions, they are important for interpretation and for future applications of this framework.

We filtered populations with uncertain windows and found stronger cue reliability for populations with more sampling years and longer sampling spans (Fig. S4). This is consistent with uncertain thermal windows being associated with limited sample size rather than necessarily pointing to the population not using temperature as a phenological cue (Bailey et al., 2022). We also conducted permutation tests and a follow-up sensitivity test based on different use of quantile-thresholds to identify robust cues. Compared to the main results based on the 0.9 quantile, the effect sizes of models based on the 0.95 quantile were identical in direction and similar in magnitude, with thermal plasticity being somewhat stronger when using the 0.95 quantile. As we found no evidence consistent with a shifting of the cue window timing, we conclude that it is unlikely that the proximate thermal cue has, over time, shifted to occur earlier or later during the season and that the overall cue windows that we identified and used in the analyses are relatively robust. However, we recognise that it is possible that this test is biased toward the null as, in the main cue window search, we screened for the period during the season that best predicts cue timing across the entire study period. Nevertheless, if the cue had shifted between the two periods, we expect that indications of this would have been evident through this sensitivity test. By design, the mean‑centering can induce statistical coupling between within‑ and between‑effects, meaning, e.g., that estimation errors in one can influence the other (Phillimore et al., 2010; Westneat et al., 2020). While our robustness checks help ensure this was not a major issue, this statistical caveat is worth nothing when interpreting these results or if applying this approach.

While the significant shift across taxa towards steeper thermal reaction norms that we found show interesting indications of contemporary evolution as a response to climate change, it is important to note that if temperature variability within short-term periods have increased through time, this would make it more likely to statistically identify steeper short‑term slopes. Such results could be misinterpreted as pointing to an evolutionary shift in reaction norm slopes. However, our test of temperature trends during the spring season showed decreasing temperature variability (cv) over time (Fig. S2), suggesting the steepening signal that we identify is not purely an artifact of variance inflation towards later decades. This potential caveat is also worth noting when replicating this approach with other data.

Our sliding window approach for identifying the most likely cue window, including population filtering, permutation tests, and the two‑period sensitivity analyses testing potential shifts in cue over time collectively support that cue windows did not shift appreciably, and that temperature during the identified cue window is a likely proximate cue for egg‑laying in the retained populations. Some species (e.g., capital breeders like the whooper swan) may use non‑local or lagged cues, such as winter severity, snowmelt timing, thermal sums, or photoperiod, and this may have caused some stochasticity or spurious species-specific results. However, the year term in the two main models, with which we tested if other environmental conditions than the identified thermal cues pointed to shifts in the reaction norms, indicated that no other temporal trends have likely given rise to the results we obtained. Instead, the approach we present here combined with the meta-analytical testing should be robust to many of the above-described caveats and offer an overall picture of in what way contemporary evolution may have acted and continue to act on breeding phenology.

Application of a sliding window approach and retention of those populations with a strong signal will, nevertheless, introduce some biases to our meta-analysis that need to be made explicit. First, our focal populations are best considered as a sample of the bird species for which breeding phenology is temperature sensitive. Second, there is some potential that the multiple testing element of the sliding window approach will over-estimate the effect size for the temperature-phenology relationship. Third, whilst the standard error of the phenology-temperature relationship would be correct if only one window had been considered, a standard error obtained from a sliding window approach will tend to be an underestimate, because it does not capture uncertainty in the slope estimate arising from uncertainty in the timing of the window.

Macphie, K. H., Morris, S., Morris, A., Guinness, F. E., Clutton-Brock, T. H., Kruuk, L. E. B., & Pemberton, J. M. (n.d.). Addressing biases in sliding window analysis gives new insight into the response of parturition date to weather in a wild mammal. *Oikos*, *n/a*(n/a), e11372. https://doi.org/10.1002/oik.11372

Phillimore, A. B., Hadfield, J. D., Jones, O. R., & Smithers, R. J. (2010). Differences in spawning date between populations of common frog reveal local adaptation. *Proceedings of the National Academy of Sciences*, *107*(18), 8292–8297. https://doi.org/10.1073/pnas.0913792107

Phillimore, A. B., Leech, D. I., Pearce-Higgins, J. W., & Hadfield, J. D. (2016). Passerines may be sufficiently plastic to track temperature-mediated shifts in optimum lay date. *Global Change Biology*, *22*(10), 3259–3272. https://doi.org/10.1111/gcb.13302

Simmonds, E. G., Cole, E. F., & Sheldon, B. C. (2019). Cue identification in phenology: A case study of the predictive performance of current statistical tools. *Journal of Animal Ecology*, *88*(9), 1428–1440. https://doi.org/10.1111/1365-2656.13038

Tansey, C. J., Hadfield, J. D., & Phillimore, A. B. (2017). Estimating the ability of plants to plastically track temperature-mediated shifts in the spring phenological optimum. *Global Change Biology*, *23*(8), 3321–3334. https://doi.org/10.1111/gcb.13624

van de Pol, M., Bailey, L. D., McLean, N., Rijsdijk, L., Lawson, C. R., & Brouwer, L. (2016). Identifying the best climatic predictors in ecology and evolution. *Methods in Ecology and Evolution*, *7*(10), 1246–1257. https://doi.org/10.1111/2041-210X.12590

Westneat, D. F., Araya-Ajoy, Y. G., Allegue, H., Class, B., Dingemanse, N., Dochtermann, N. A., Garamszegi, L. Z., Martin, J. G. A., Nakagawa, S., Réale, D., & Schielzeth, H. (2020). Collision between biological process and statistical analysis revealed by mean centring. *Journal of Animal Ecology*, *89*(12), 2813–2824. https://doi.org/10.1111/1365-2656.13360
